# A tissue-specific ubiquitin switch coordinates brain, craniofacial, and skin development

**DOI:** 10.1101/2022.09.26.509591

**Authors:** Anthony J. Asmar, Rita M. Yazejian, Youmei Wu, Jason C. Collins, Jenny Hsin, Jean Cho, Andrew D. Doyle, Samhitha Cinthala, Marleen Simon, Richard H. van Jaarsveld, David B. Beck, Laura Kerosuo, Achim Werner

## Abstract

The molecular mechanisms that coordinate patterning of the embryonic ectoderm into spatially distinct lineages to form the nervous system, epidermis, and craniofacial structures are unclear. Here, biochemical disease-variant profiling reveals a posttranslational pathway that drives early ectodermal differentiation in the vertebrate head. The anteriorly expressed ubiquitin ligase CRL3-KLHL4 restricts signaling of the ubiquitous cytoskeletal regulator CDC42. The major substrate of CRL3-KLHL4 is the canonical CDC42 effector kinase PAK1 that monoubiquitylation switches into a CDC42 inhibitor. Loss of CRL3-KLHL4 or a disease-associated *KLHL4* variant reduce PAK1 ubiquitylation causing overactivation of CDC42 signaling and defective ectodermal patterning and neurulation. Thus, tissue-specific, ubiquitin-dependent restriction of CDC42 signaling is essential for face, brain, and skin formation, demonstrating how cell-fate and morphometric changes are coordinated for faithful organ development.

---

During early development, the ectoderm layer of vertebrate embryos undergoes neurulation, involving drastic morphological changes to form the neural tube, while it is simultaneously patterned into three distinct domains. The lateral non-neural ectoderm domain will form the protective epidermis and the medial neural plate domain gives rise to the central nervous system (CNS) that forms the brain in the head and the spinal cord in the posterior body. In between these two domains is the neural plate border, which forms the cranial placodes and the neural crest cells that will generate multiple derivatives, including the peripheral nervous system and, in the head region, the craniofacial skeleton^1^. A failure to faithfully establish and differentiate ectodermal domains results in a range of congenital birth defects affecting the development of the central and the peripheral nervous system, craniofacial structures, and the skin^2-5^. Extensive studies in different model systems have identified key morphogens gradients in the embryo that induce major transcriptional changes driving ectodermal patterning into the respective domains^6-17^. Yet, molecular mechanisms that determine these initial cell-fate decisions and coordinate them with morphometric cell changes to spatially separate domains within the neurulating ectoderm have remained elusive^18^.

Cullin-RING E3 ubiquitin ligases (CRLs) are a large class of ∼300 modular enzymes that control critical aspects of human development and physiology ^19-22^. The CRL3 sub-family of these enzymes consist of a stable catalytic core complex, CUL3-RBX1, which is paired with one of ∼120 interchangeable substrate adaptors that contain a BTB domain^23-25^. While the BTB domain connects the adaptor to CUL3-RBX1, other dedicated domains in the adaptor (e.g. KELCH repeats) recognize specific substrates (***Fig. 1A***). The assembly and disassembly of particular CRL3-substrate complexes is tightly regulated within cells and involves reversible modification with NEDD8 (neddylation), the substrate adaptor exchange factor CAND1^26-32^, and co-adaptors to help recruit substrates and to coordinate the assembly process^33-35^. CRL3s often catalyze non-degradative monoubiquitylation^22, 34-37^ to control organismal development and tissue homeostasis^38-40^. Specifically, CRL3s play crucial roles in ectodermal differentiation, as evidenced by the fact that more than 20 different loss-of-function variants in the *CUL3* locus are associated with neurodevelopmental and craniofacial diseases^41^ (***Table S1***). However, how particular CRL3 complexes control early stages of ectodermal patterning and neural tube formation remains unknown.

**Figure 1:**
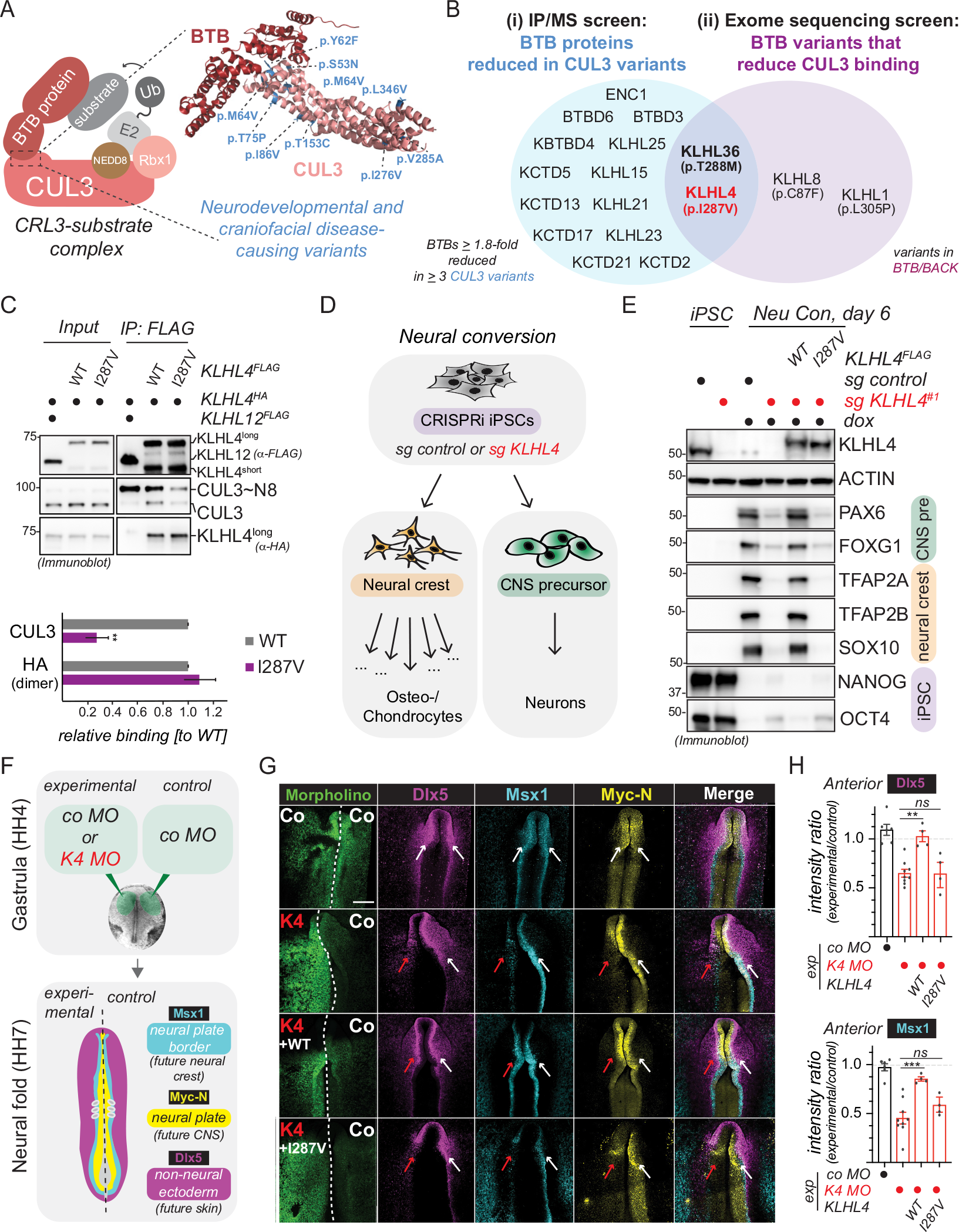
CRL3-KLHL4 ubiquitylation activity controls ectodermal patterning and neurulation in the vertebrate head. **A**) Model of a substrate-engaged CRL3 complex with an example of a crystal structure of a CUL3-BTB interface (pdb: 4APF) highlighting the positions of missense variants in CUL3 that have been shown to cause neurodevelopment and craniofacial phenotypes in patients. We have used these mutations as a tool in our mass spectrometry (MS) screen. **B**) Schematic overview of the IP/MS screen (i) and the exome sequencing screen (ii) for novel CRL3-BTB complexes with functions during ectodermal differentiation, identifying CRL3-KLHL4 and CRL3-KLHL36 as hits. **C**) The KLHL4 p.I287V patient variant does not impact KLHL4 dimerization, but reduces interaction with CUL3, as shown by immunoblot analysis of FLAG-IP fractions from HEK293T cells. Note that ectopically expressed KLHL4 is present in two isoforms (short and long), likely originating from differential usage of start codons at M1 and M129, respectively (***Fig. S2N***). Graph depicts quantification of the anti-FLAG IP experiments (n=3 biological replicates, error bars = s.d., ** = p < 0.01, student t-test). **D**) Schematic overview of neural conversion, a differentiation paradigm that directs human pluripotent stem cells towards the indicated ectodermal cell fates. CNS = central nervous system. CRL3-KLHL4 ubiquitylation activity is required for neural conversion. Control or KLHL4-depleted CRISPRi iPSCs stably expressing sgRNA-resistant and doxycycline-inducible wildtype (WT) or patient variant (I287V) KLHL4^FLAG^ were generated followed by treatment with doxycycline (dox) and neural conversion for 6 days as indicated. Differentiation was monitored by immunoblotting using indicated antibodies against specific lineage markers. ACTIN = loading control. Bilateral injections of fluorescein-labeled translation blocking morpholinos were electroporated into the ectoderm at gastrula stage (HH4) and incubated until the embryos reached the neural folds stage (HH7-8) and analyzed. **G**) Loss of KLHL4 (K4) reduces anterior expression of markers for all ectodermal domains, including non-neural ectoderm (Dlx5), neural plate border (Msx1), and neural plate (Myc-N), whereas control embryos show no difference between sides as shown by whole mount HCR fluorescent in situ hybridization. Morpholino-induced loss of KLHL4 is is rescued by co-expression of WT KLHL4, but not with the patient mutation KLHL4 I287V. **H**) Quantification of fluorescence intensities of Dlx5 and Msx1 show statistically significant rescue of KLHL4 MO with WT KLHL4, but not with patient mutation I287V KLHL4 (Dlx5: n=4-9 embryos as indicated, ** = p < 0.01, one-way ANOVA; Msx1: n=4-9 embryos as indicated, *** = p < 0.01, one-way ANOVA).

Here, by combining systematic human disease variant screening with biochemical, iPSC, and chick embryo approaches, we identify CUL3 in complex with its substrate adaptor KLHL4 (CRL3-KLHL4) as central to a novel posttranslational pathway that drives neurulation and ectodermal domain formation. We find that monoubiquitylation by CRL3-KLHL4 catalyzes an effector-to-inhibitor conversion to restrict cytoskeletal signaling of the small GTPase CDC42 in the developing vertebrate head. Our data thus uncover a previously unrecognized principle of small GTPase signaling that coordinates cell-fate and morphometric changes to establish the future skin, brain, and craniofacial skeleton and, upon dysregulation, contributes to craniofacial and brain malformations.

## RESULTS

### CRL3-KLHL4 ubiquitylation activity controls ectodermal patterning and neurulation

To identify CRL3s with previously unrecognized functions during ectodermal differentiation, we biochemically profiled known or candidate neurodevelopmental and craniofacial disease-causing variants in the CUL3-BTB interface (***Fig. 1A***). We identified KLHL4 and KLHL36 as the only BTB adaptors that were both (i) reduced in immunoprecipitation (IP) fractions of several disease-associated CUL3 variants and (ii) for which we found variants in patients with undiagnosed developmental diseases that reduced their interaction with CUL3 (***Fig. 1B, Fig. S1, Fig. S2, Table S2, S3***). Given its previous connection to craniofacial development^42, 43^, we focused on *KLHL4* for follow-up studies. The maternally inherited hemizygous missense variant in KLHL4 (NM_019117.4: c.859A>G [p.(Ile287Val)] that we identified in our exome sequencing screen is present in a male proband, exhibiting severe intellectual disability and multiple congenital anomalies, including brain malformations (for a more detailed patient description refer to ***Note S1***). The KLHL4 p.I287V variant significantly reduced binding to unmodified and neddylated CUL3, while not affecting KLHL4 dimerization (***Fig. 1C, Fig. S2K***).

To test for a function of CRL3-KLHL4 during ectodermal differentiation, we generated control and KLHL4-depleted CRIPSRi iPSCs^44^ and subjected these cell lines to neural conversion^45^ (***Fig. 1D***). Utilizing self-made monoclonal antibodies against KLHL4, we observed a marked reduction of KLHL4 protein levels during neural conversion of control iPSCs (***Fig. 1E***), raising the possibility that KLHL4 could play a role during early stages of ectodermal differentiation. Indeed, while we observed no substantial differences in cell growth or the expression of pluripotency markers OCT4 and NANOG in the ground state, there was a notable defect to form neural and neural crest derivatives when comparing control and KLHL4-depleted iPSCs (***Fig. 1E, Fig. S2O***). This was apparent by the loss of neural crest markers (including SOX10 and TFAP2A/B), reduced expression of CNS markers (including the forebrain marker FOXG1 and the neural stem cell marker PAX6), and failure to downregulate OCT4 and NANOG, as evidenced by immunoblotting, quantitative polymerase chain reaction (qPCR), and immunofluorescence (***Fig. 1E, Fig. S3A,B***). We could further corroborate these findings during an alternative *in vitro* protocol (***Fig. S4***). Importantly, these effects could be rescued by wildtype (WT) KLHL4, but not by the CUL3-binding-deficient KLHL4 p.I287V patient variant (***Fig. 1E, Fig. S3A,B***), demonstrating that the ubiquitylation activity of CRL3-KLHL4 is required to support ectodermal differentiation of pluripotent stem cells.

To corroborate the above findings *in vivo*, we next performed experiments in chick embryos, an amniote model with high resemblance to human development^46^. In situ hybridization analysis revealed *Klhl4* to be exclusively expressed in the ectoderm and restricted to the anterior part of the developing embryo that will form the head (***Fig. S5A***), suggesting a tissue-specific role of CRL3-KLHL4 during ectodermal cell-fate specification. Indeed, using translation-blocking morpholinos against *KLHL4* (K4 MO) (***Fig. 1F***), we found that loss of KLHL4 severely interfered with ectodermal lineage commitment of all respective domains (***Fig. 1G, Fig. S5B***). This was evident in whole embryos, in which expression of the neural plate/CNS marker *MycN*, the neural plate border marker *Msx1*, and the non-neural ectoderm marker *Dlx5*, were significantly reduced in the KLHL4 MO-injected side as compared to the side with the control MO as shown by multichannel fluorescent *in situ* hybridization (***Fig. 1G,H Fig. S5B***). Consistent with the expression pattern of KLHL4, these phenotypes were only present in the anterior, but not in the posterior part of the embryo (***Fig. S5B-D***). Furthermore, analysis of corresponding anterior transverse sections revealed that depletion of KLHL4 prevented proper neurulation as shown by significantly less folding of the neural plate as compared to the control side (***Fig. S5E,F***). The sections also revealed that loss of KLHL4 significantly compromised the establishment of defined borders between the distinct ectodermal domains, measured as increase in the overlap of the expression of the markers representing the distinct domains (***Fig. S5G***). Consistently, we detected a significant increase in the length of the neural plate border domain upon KLHL4 depletion when measured by immunostaining the neural plate borders with a Pax7 antibody (***Fig. S5H,I***). Importantly, the KLHL4 MO knockdown phenotype was rescued with WT KLHL4, but not with the CUL3-binding-deficient KLHL4 p.I287V patient variant, which continued to show an aberrant ectodermal patterning phenotype (***Figure 1G,H)***. Taken together, these results suggest that during the early neurulation stage of embryo development, tissue-specific ubiquitylation by CRL3-KLHL4 ensures neural plate folding and cell-fate specification of the head ectoderm into functionally distinct domains, and impairment of this activity through mutations likely contributes to neurodevelopmental and craniofacial diseases.

### CRL3-KLHL4 uses GIT-PIX complexes as co-adaptors to recruit and multi-monoubiquitylate PAK1

To elucidate the molecular mechanisms by which CRL3-KLHL4 allows proper distinction of ectodermal domains and neurulation, we employed an approach that had previously allowed us to define stem cell-related signaling pathways^35, 47^. We used compPASS mass spectrometry^48, 49^ to identify candidate substrates of CRL3-KLHL4. We determined proteins that bind to WT KLHL4, KLHL4 p.I287V patient variant, and a more severe CUL3-binding mutant (KLHL4ΔCUL3). These interaction networks revealed GIT1, GIT2, α-PIX, β-PIX, and PAK1 as predominant interactors that bound equally to WT KLHL4, KLHL4 p.I287V, and KLHL4 ΔCUL3, indicating interaction with the substrate-binding domain of KLHL4 (***Fig. 2A***). These proteins are components of GIT-PIX-PAK assemblies, which regulate cytoskeletal signaling by modulating the activity of the small GTPases CDC42 and RAC1^50^. We confirmed the CRL3-KLHL4 interaction with the GIT-PIX-PAK signaling module by IP/immunoblotting (***Fig. S6A***) and showed that the same associations occur at the endogenous level in hESCs (***Fig. 2B***).

**Figure 2:**
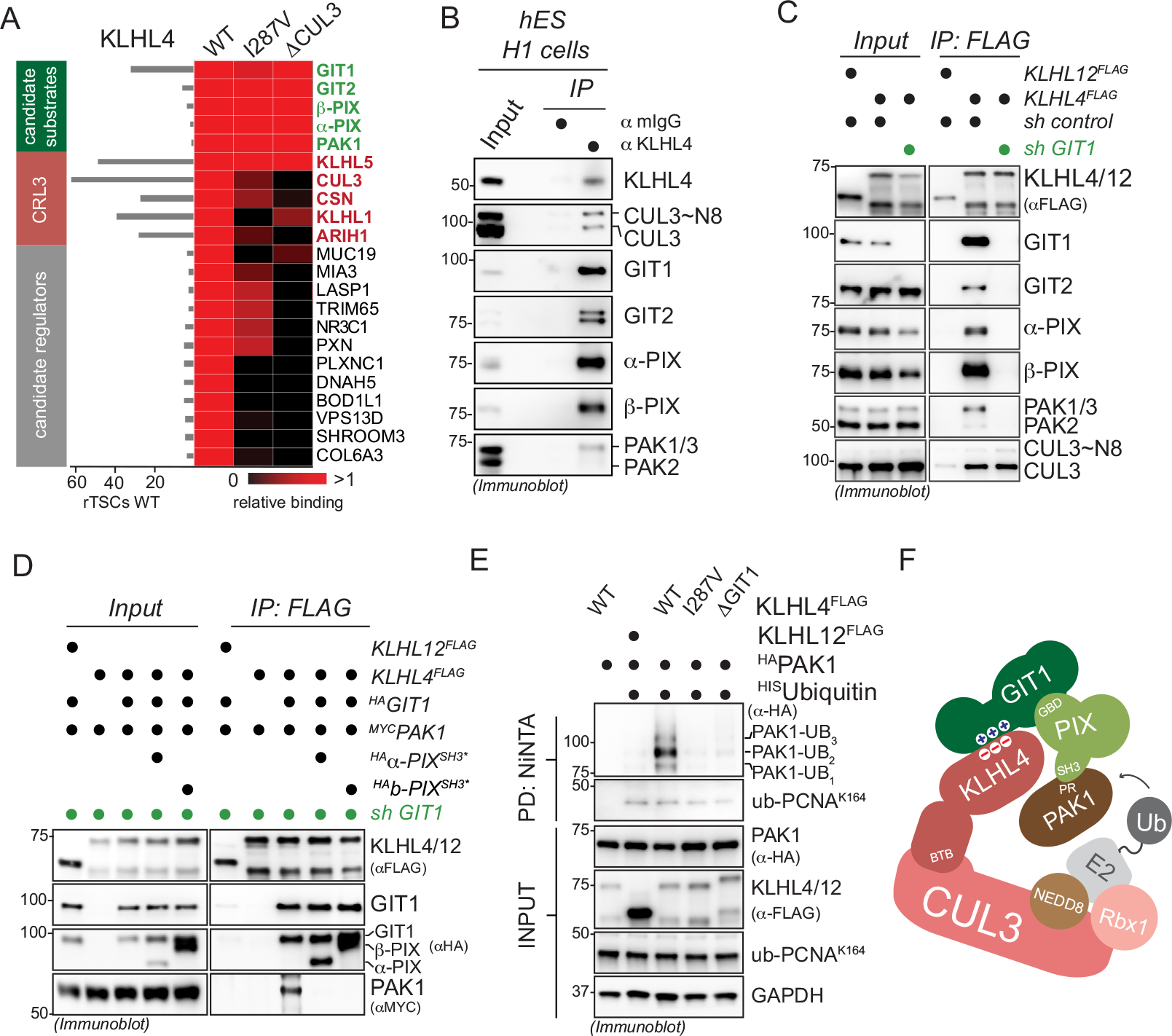
CRL3-KLHL4 uses GIT1-PIX complexes as co-adaptors to recruit and multi-monoubiquitylate PAK1. **A**) Components of the GIT-PIX-PAK signaling module are candidate substrates of CRL3-KLHL4, as suggested by their ability to bind to wild-type and CUL3-binding-deficient KLHL4. FLAG-tagged wild-type KLHL4 (WT), patient variant KLHL4 (I287V), and a more severe CUL3-binding mutant of KLHL4 (ΔCUL3) were analyzed by compPASS-based mass spectrometry. The heatmap depicts the relative binding of interactors identified for KLHL4 WT in the respective KLHL4 variant (black = no interaction, red = equal or more interaction). A quantification of normalized total spectral counts (TSCs) of three biological replicates is shown for KLHL4 WT. **B**) Endogenous KLHL4 interacts with the GIT-PIX-PAK signaling module in hESCs (H1 line), as evidence by immunoblot analysis of endogenous anti-KLHL4 IPs. mIgGs were used as IP control. **C**) CRL3-KLHL4 engages the GIT-PIX-PAK signaling module through GIT1, as evidenced by immunoblot analysis of anti-FLAG IPs from control or GIT1-depleted HEK293T cells. **D**) GIT-PIX complexes recruit PAK1 to KLHL4. GIT1-depleted HEK 293T cells were co-transfected with indicated combinations of KLHL4^FLAG^ of KLHL12 ^FLAG^, ^MYC^PAK1, shRNA-resistant GIT1^HA^, and PAK1-binding-deficient α/α-PIX mutants (^HA^ α/α-PIX ^SH3*^), lysed, and subjected to anti-FLAG IP followed by immunoblotting with indicated antibodies. Of note, KLHL4 can only bind PAK1 when shRNA-resistant GIT1 is expressed, and this association can be blocked by co-expression of PAK-binding-deficient PIX proteins. **E**) CRL3-KLHL4 ubiquitylates PAK1 in cells. Ubiquitylated proteins were purified under denaturing conditions from RPE-1 cells expressing ^HIS^ubiquitin and denoted combinations of ^HA^PAK1, KLHL12^FLAG^, and KLHL4^FLAG^ variants. Modified proteins were detected by immunoblotting using the indicated antibodies. PCNA is ubiquitylated by a distinct E3 ligase and hence is used as a control for general ubiquitylation efficiency. **F**) Cartoon depicting a simplified model on how CRL3-KLHL4 uses GIT1-PIX complexes as co-adaptors to bind and ubiquitylate PAK1.

To determine how CRL3-KLHL4 engages the GIT-PIX-PAK signaling module, we performed a series of biochemical and cell-based experiments. IP of KLHL4 from GIT1-depleted cells showed that GIT1 was required for KLHL4 to recognize GIT2, α-PIX, β-PIX, PAK1, or PAK3, but not CUL3 (***Fig. 2C, Fig. S6B***). Truncation and mutational analyses revealed three positively charged amino acids (R564, R566, and K567) in GIT1 that, when substituted with negatively charged amino acids (GIT1^RK3D^), impaired interaction with KLHL4 (***Fig. S7A-C***). Only R564 is present in GIT2, the otherwise highly conserved homologue of GIT1 (***Fig. S7D***), providing a molecular rationale for why we find only GIT1, and not GIT2, required for targeting the GIT-PIX-PAK module to CRL3-KLHL4 (***Fig. S7E***). Through structural modeling, we identified a complementary negatively charged surface on the bottom of the KLHL4 β-propeller that when disrupted by charge swap mutations (D475R,D544R,E546R,D591R) reduced binding to the GIT-PIX-PAK module (KLHL4ΔGIT1, ***Fig. S8A,B, S6B***), indicating that GIT1 binds KLHL4 on the opposite side of where substrate binding usually occurs in BTB-KELCH proteins^51-53^. Indeed, using recombinantly purified proteins, we were able to *in vitro* reconstitute binding of KLHL4 to GIT1, but not to GIT1^RK3D^ (***Fig. S8C***). Collectively, our results show that CRL3-KLHL4 engages the GIT-PIX-PAK signaling module through a direct electrostatic interaction with GIT1.

Since our results revealed that the GIT1-KLHL4 binding interface is distal from the canonical substrate binding site, we reasoned that this interaction might serve to recruit PIX and/or PAK proteins to the CRL3-KLHL4 complex for ubiquitin ligation. Indeed, building on previously mapped interactions of GIT-PIX-PAK module^50, 54^ (***Fig. S9A***), we found that ectopically expressed GIT1 increased association of endogenous α-PIX and β-PIX to affinity-purified KLHL4^FLAG^, while KLHL4 binding-deficient GIT1^Y563D^ (***Fig. S7C***) sequestered these proteins and prevented their association with KLHL4 (***Fig. S9B***). In addition, IP experiments from GIT1-depleted cells revealed that binding of PAK1 to KLHL4 was only supported upon rescue by expressing shRNA-resistant GIT1, and this association could be blocked by co-expression of PIX proteins (α-PIX^SH3*^, α-PIX ^SH3* 55^) that are unable to bind PAK1 (***Fig. 2D***). Thus, engagement of CRL3-KLHL4 to the GIT-PIX-PAK signaling module occurs through GIT1 that binds to PIX proteins, which can further recruit PAK1 (***Fig. S9C***).

To test whether the above interactions result in ubiquitylation, we purified ubiquitin conjugates from cells under denaturing conditions. Intriguingly, we found that KLHL4, but not the CUL3-binding deficient KLHL4 p.I287V patient variant, induced robust ubiquitylation of PAK1 (***Fig. 2E***), while under similar experimental conditions GIT1-β-PIX complexes were not ubiquitylated in a KLHL4-dependent manner (***Fig. S10A***). PAK1 ubiquitylation was not supported by KLHL4 ΔGIT1, indicating that CRL3-KLHL4 requires GIT1-PIX complexes to form efficient ubiquitin ligation assemblies with PAK1 (***Fig. 2E***). Corroborating this notion, a PAK1 mutant unable to interact with PIX proteins (PAK1^ΔPIX^) was also deficient in KLHL4-dependent ubiquitylation (***Fig. S10B,C***). KLHL4-dependent PAK1 modifications are likely multi-monoubiquitylation events, as we predominantly observed attachment of up to three ubiquitin molecules (***Fig. 2E, Fig. S10C***) and a lysine-less version of ubiquitin still supported the same pattern of modification for PAK1 (***Fig. S10D***). Consistent with such multi-monoubiquitylation, which typically does not mediate proteasomal degradation, we did not detect changes in the steady state levels of PAK1 or other components of the GIT-PIX-PAK module upon loss of KLHL4 in cells (***Fig. S10E***). From these results, we infer that CRL3-KLHL4 utilizes GIT-PIX complexes as co-adaptors to recruit and multi-monoubiquitylate PAK1 (***Fig. 2F***).

### CRL3-KLHL4 regulates ectodermal differentiation through monoubiquitylation of PAK1

To verify that PAK1 ubiquitylation is important for the function of CRL3-KLHL4 in ectodermal development, we performed a series of iPSC differentiation experiments. First, similar to loss of KLHL4, iPSCs expressing only GIT-PIX-PAK-binding deficient KLHL4ΔGIT1 failed to support CNS precursor and neural crest cell formation (***Fig. S11A-C***). Second, the same aberrant neural conversion program was observed if we depleted GIT1, α−PIX, and PAK1/2/3, while depletion of KLHL5, GIT2, or α−PIX caused less dramatic changes (***Fig. S11D,E***). These result are consistent with our findings that GIT1, but not GIT2, is required for PAK1 recruitment to KLHL4 (***Fig. S7E***) and further suggest that during differentiation it is mainly GIT1 - β−PIX complexes that bridge interactions of CRL3-KLHL4 with PAK1 for ubiquitin ligation. Third, we found that constitutively ubiquitylated PAK1, as mimicked by ubiquitin fused to PAK1^33, 56, 57^ (^HA^PAK1-Ub), was able to restore the neural conversion program of KLHL4-depleted iPSCs, while this was not the case for its unmodified counterpart ^HA^PAK1 (***Fig. 3A,B***). In addition, PAK1 fused to ubiquitin mutated in its hydrophobic patch (^HA^PAK1-Ub^I44A^) was markedly reduced in its ability to rescue differentiation, suggesting that monoubiquitylated PAK1 mediates cell-fate determination at least in part through recognition by a ubiquitin-binding effector protein. Strikingly, constitutively ubiquitylated PAK1-Ub also partially rescued KLHL4-depletion-induced defects in ectodermal patterning during chick development in a dose-dependent manner, as quantified by the expression of the neural plate border marker *Msx1* (***Fig. 3C,D, Fig. S12A***). In contrast, non-ubiquitylated PAK1 was not able to restore normal ectodermal cell-fate specification in KLHL4 MO-injected embryos and even resulted in excessive and ectopic expression of *Msx1* in the extra-embryonic tissue in some cases (***Fig. S12A***). Collectively, these results suggest that GIT1-β−PIX-dependent mono-ubiquitylation of PAK1 is a central mechanism underlying the function of CRL3-KLHL4 in controlling ectodermal patterning (***Fig. S12B***). Consistent with our findings, genetic lesions in the GIT-PIX-PAK signaling axis are known to cause congenital diseases affecting brain and craniofacial development with phenotypic overlap to *CUL3* and *KLHL4* patients^58-61^.

**Figure 3:**
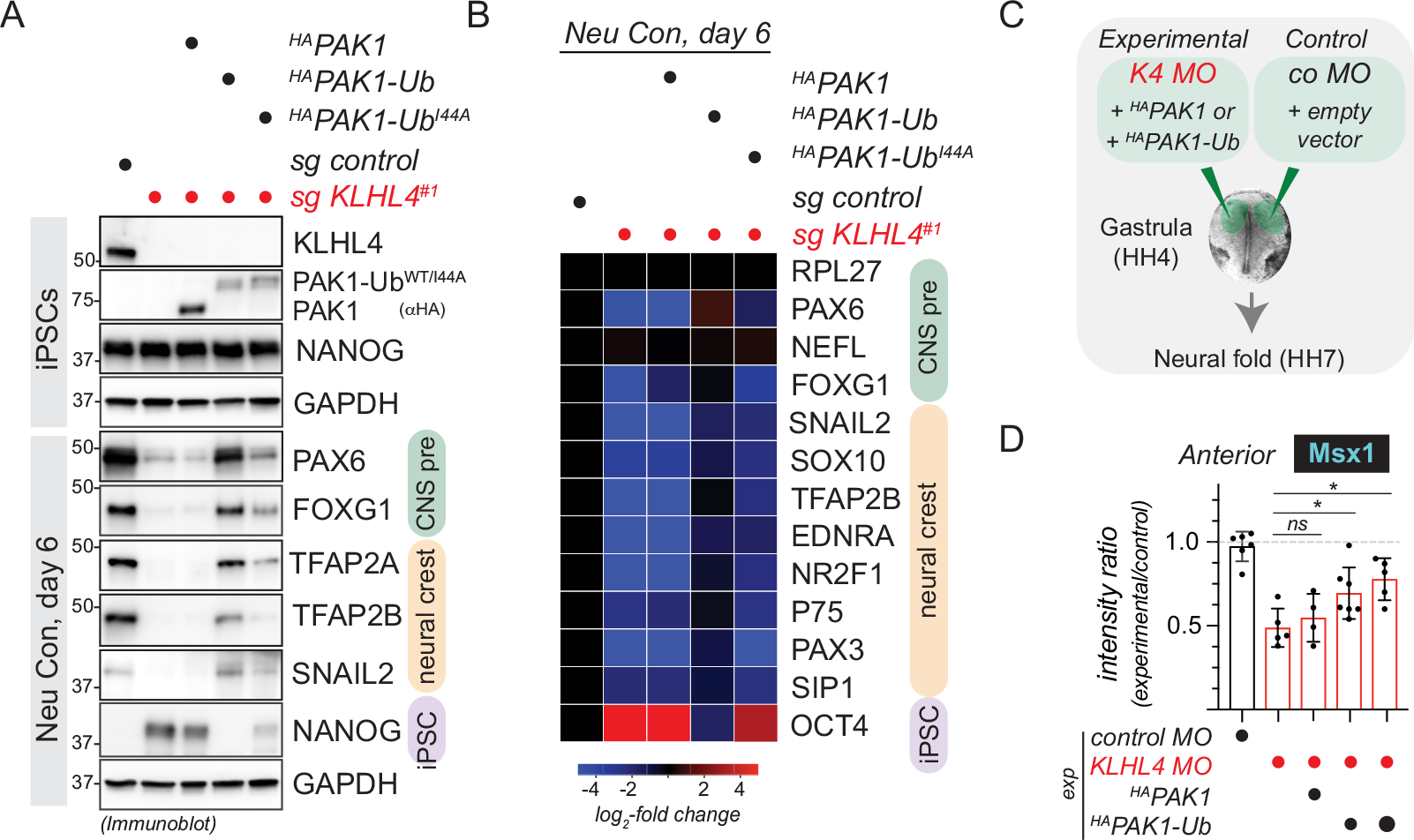
Monoubiquitylation of PAK1 controls ectodermal differentiation. **A)** Ubiquitylated PAK1 can rescue neural conversion of KLHL4-deficient iPSCs. Control or KLHL4-depleted CRISPRi iPSCs were reconstituted with doxycycline-inducible wildtype ^HA^PAK1, ^HA^PAK1-Ub, or ^HA^PAK1-Ub^I44A^, treated with doxycycline (dox) and subjected to neural conversion for 6 days. Differentiation was monitored by immunoblotting using antibodies against indicated lineage markers. **B**) Ubiquitylated PAK1 can rescue neural conversion of KLHL4-deficient iPSCs, as monitored by qPCR analysis of iPSCs treated and differentiated as in panel CNS precursor markers = green, neural crest markers = orange, and iPSC markers = purple. Marker expression was normalized to control iPSCs and RPL27 was used as endogenous control. Heatmap depicts the average of 2 biological replicates with 3 technical replicates each. **C**) Experimental design for *in vivo* electroporations testing whether constitutively ubiquitylated PAK1 can rescue KLHL4 depletion. **D**) Morpholino-induced loss of KLHL4 phenotype is rescued in a dose-dependent manner by co-expression of ubiquitylated PAK1, but not by the non-ubiquitylated WT PAK, as quantified by ratio of Msx1 fluorescence intensity between experimental and control side from embryos stained with whole mount HCR FISH, which show significant difference between KLHL4 MO and PAK1-Ub rescue. For reference, Msx1 intensity rations of embryos that were injected with control MO on both sides are shown. n = 4-7 embryos

### Monoubiquitylation converts PAK1 into an inhibitor of CDC42 signaling

We next aimed at determining how monoubiquitylation of PAK1 regulates cellular signaling to ensure proper ectodermal patterning. The GIT-PIX-PAK axis participates in remodeling of the actin cytoskeleton through Rho-family small GTPases^50, 62^ and, consistent with these studies, we found CRL3-KLHL4 to regulate cell shape and actin dynamics (***Fig. S13A-K, Movie S1***). We therefore hypothesized that CRL3-KLHL4-dependent mono-ubiquitylation of PAK1 might regulate RhoA, RAC1, or CDC42 signaling to ensure ectodermal cell-fate commitment. Intriguingly, using affinity pull down of GTP-bound GTPases from cells, we found that KLHL4 depletion significantly increased CDC42 activity in iPSC undergoing early stages of neural conversion, while RhoA or RAC1 were not affected (***Fig. 4A,B***). This effect could be rescued by ectopic expression of WT KLHL4, but not by the CUL3-binding- or GIT-PIX-PAK-binding-deficient variants of KLHL4 (I287V and ΔGIT1, respectively) (***Fig. S14A,B***), suggesting that CRL3-KLHL4-dependent ubiquitylation of PAK1 is a means to restrict CDC42 activity. Indeed, normal levels of CDC42 activation in KLHL4-depleted cells could be restored by expression of constitutively ubiquitylated PAK1 (^HA^PAK1-Ub), but not its unmodified counterpart (^HA^PAK1) and only partially by PAK1 fused to the ubiquitin hydrophobic patch mutant (^HA^PAK1-Ub^I44A^) (***Fig. 4C, Fig. S14C***). Similarly, co-depletion of the GTPase exchange factors (GEFs) α−PIX and β−PIX in KLHL4-deficient cells, re-established CDC42-GTP levels to that of control cells (***Fig. S14D,E***). From these experiments we conclude that during ectodermal differentiation, ubiquitylated PAK1, likely through interaction with an ubiquitin-binding effector protein, restricts CDC42 activity by counteracting *α/α−*PIX-mediated GDP-to-GTP exchange (***Fig. 4D***), revealing that monoubiquitylation can switch a canonical CDC42 effector^63, 64^ into an inhibitor.

**Figure 4:**
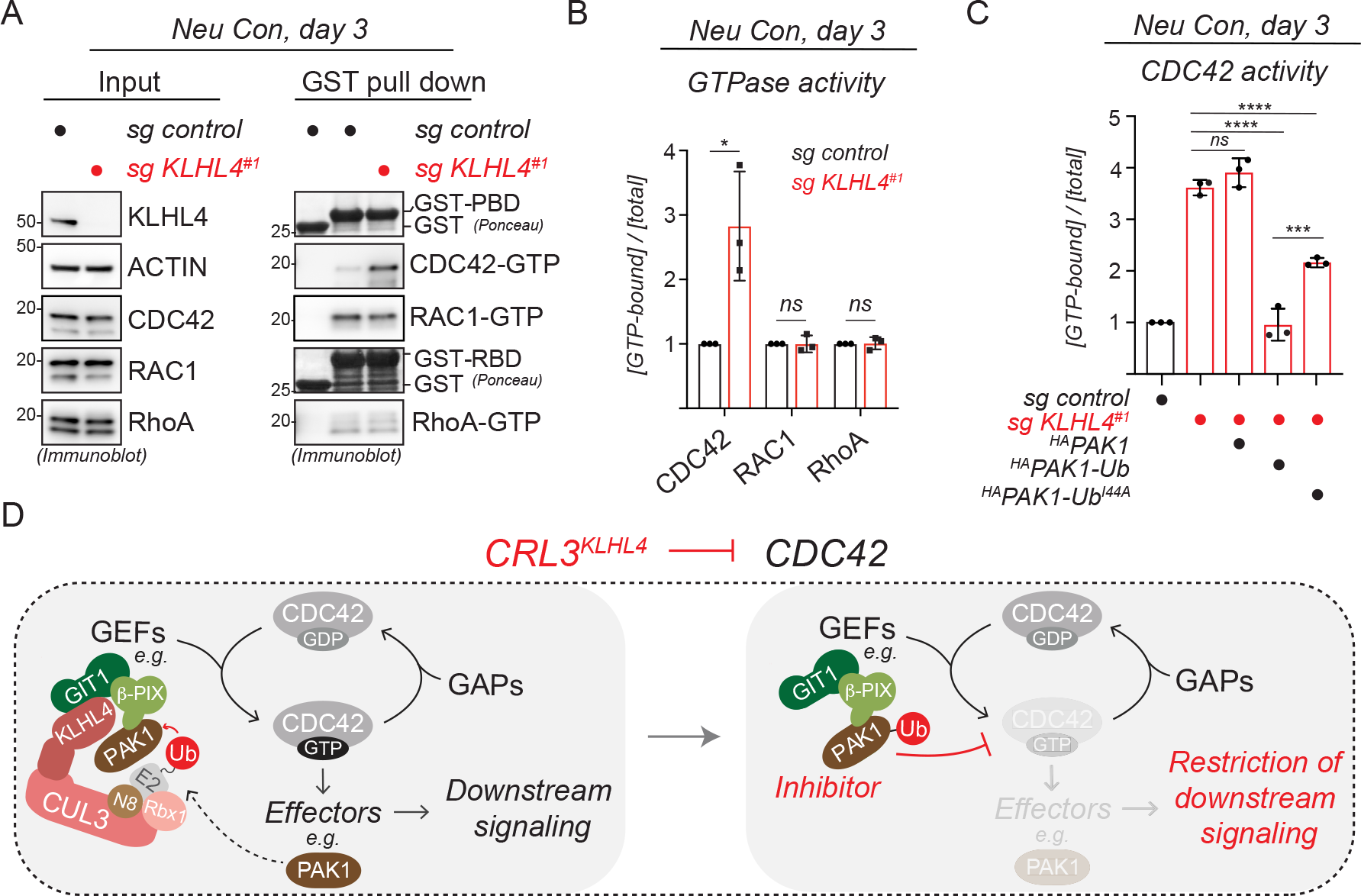
Monoubiquitylation of PAK1 restricts CDC42 signaling. **A**) KLHL4 depletion increases CDC42 activity in cells undergoing early stages of neural conversion. Control or KLHL4-depleted CRISPRi iPSCs were subjected to neural conversion for 3d, lysed, and endogenous active (GTP-bound) small GTPases were affinity-purified using GST-PBD (CDC42/RAC1) or GST-RBD (RhoA) followed by immunoblotting with indicated antibodies. GST was used as specificity control for binding. **B**) Quantification of the experiment depicted in panel A. Relative small GTPase activity was calculated by dividing the GTP-bound state to the total levels of each small GTPase followed by normalization to control cells. n = 3 biological replicates, error bars = s.d., * = p < 0.05, student t-test. **C**) Ubiquitylated PAK1 can restore normal levels of CDC42 activity in KLHL4-deficient cells undergoing early stages of neural conversion. Control or KLHL4-depleted CRISPRi iPSCs were reconstituted with dox-inducible ^HA^PAK1, ^HA^PAK1-Ub, or ^HA^PAK1-Ub^I44A^, treated with dox and subjected to neural conversion for 3d. GTP-bound CDC42 was affinity-purified and relative CDC42 activity was calculated as described above. n = 3 biological replicates, error bars denote s.d., *** = p < 0.001, **** = p < 0.001, one-way ANOVA. **D**) Model illustrating how CRL3-KLHL4 restricts CDC42 signaling during ectodermal differentiation. CRL3-KLHL4 uses GIT1-β−PIX complexes to recruit and ubiquitylate PAK1, which converts this canonical CDC42 effector into a CDC42 inhibitor that counteracts α/β−PIX-mediated CDC42 activation.

### Monoubiquitylation of PAK1 balances anterior CDC42 signaling to coordinate ectodermal patterning and neurulation

The above results suggested that loss of CRL3-KLHL4-dependent PAK1 ubiquitylation results in abnormally increased CDC42 activation. We therefore tested whether the CDC42 small molecule inhibitor, ML141 (CDC42i)^65^ would ameliorate ectodermal differentiation phenotypes in iPSCs lacking CRL3-KLHL4 activity. Intriguingly, while KLHL4-depleted iPSCs failed to undergo efficient neural conversion when cultured in presence of the vehicle control DMSO, addition of ML141 to the media rescued the aberrant differentiation program in a dose-dependent manner (***Fig. 5A,B, Fig. S15A***). To expand these observations to our *in vivo* model, we next asked whether a constitutively inactive version of CDC42 still capable of GEF binding (CDC42^T17N^) could “sponge” excessive GEF activity and thus rescue KLHL4-depletion-induced phenotypes in the embryo (***Fig. 5C***). Strikingly, co-injection of low concentrations of a plasmid encoding CDC42^T17N^ with KLHL4 MO re-established ectodermal domain specification and neural tube formation to similar levels of control MO-injected embryos (***Fig. 5D,E***). Surprisingly, in these experiments we noted that increasing the concentration of CDC42^T17N^ resulted in an over-advanced neural plate border with significantly increased levels of *Msx1* compared to the control side of the embryo, and the whole mount images indicated an acceleration of the neural plate folding (***Fig. 5D,E***). This suggests that expression of CDC42^T17N^ above a certain threshold reduces CDC42 activity below physiological levels, again resulting in aberrant formation of ectodermal domains. To further substantiate this notion, we next performed CDC42^T17N^ injections in embryos without KLHL4 MO knockdown (***Fig. S15B***). Also under these conditions, CDC42^T17N^ expression caused a more defined neural plate border and increased *Msx1* expression as compared to the control side of the embryo, indicating an acceleration of the ectodermal patterning process (***Fig. S15C,D***). Taken together, we conclude that post gastrulation, CRL3-KLHL4-dependent ubiquitylation of PAK1 in the anterior ectoderm is required to establish a precise level of CDC42 signaling. This ubiquitin-dependent regulation coordinates faithful ectodermal patterning and neural tube formation and both, too much or too little anterior CDC42 activity, results in aberrant ectodermal development.

**Figure 5:**
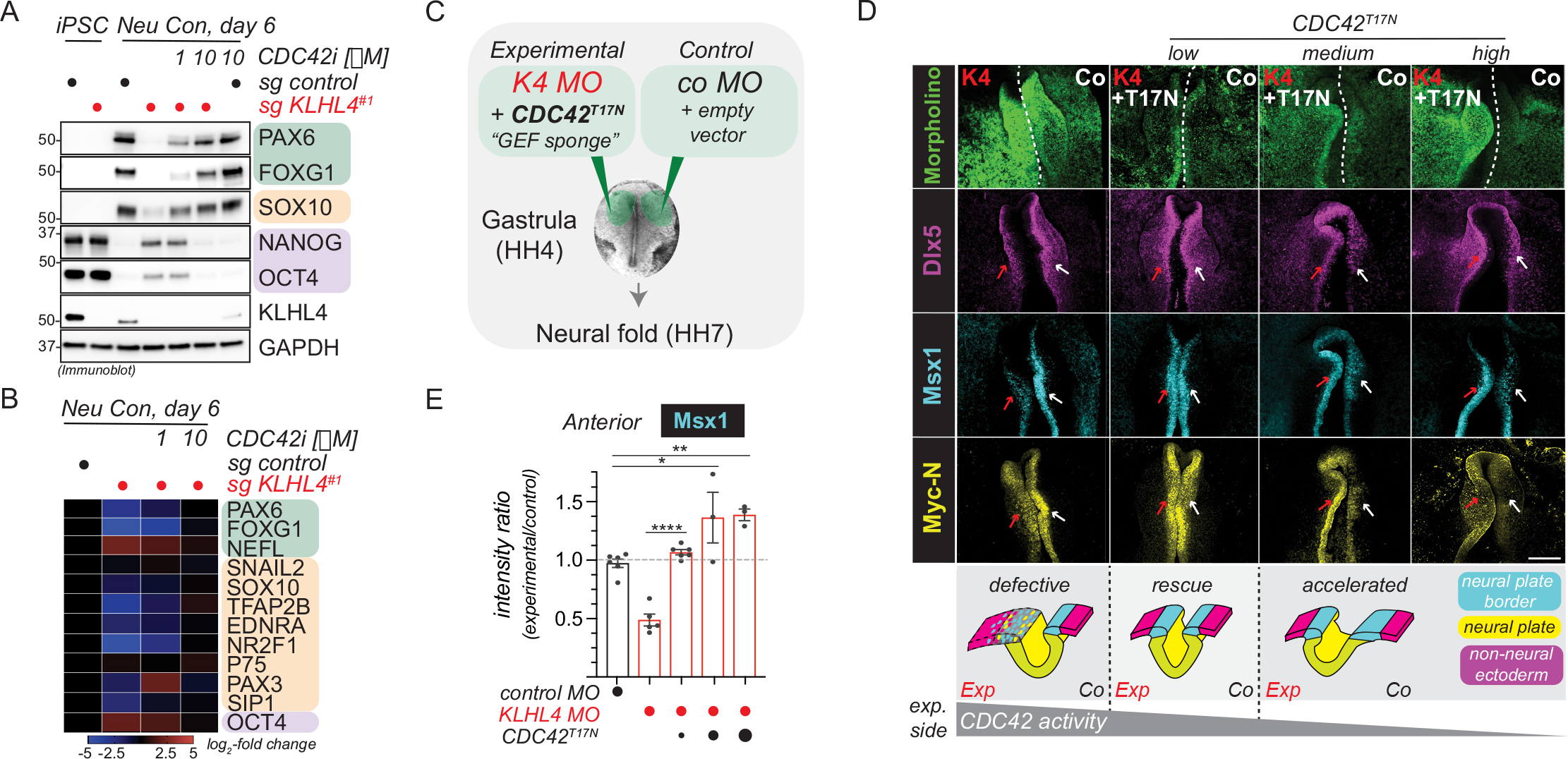
CRL3-KLHL4 balances CDC42 signaling to coordinate ectodermal patterning and neurulation in the vertebrate head. **A**) CDC42 inhibition rescues neural conversion of KLHL4-deficient iPSCs. Control or KLHL4-depleted CRISPRi iPSCs were treated with DMSO or indicated concentrations of the CDC42 inhibitor ML141, subjected to neural conversion for 6d, and analyzed by immunoblot analysis for expression of CNS precursor markers (green), neural crest markers (orange), and pluripotency markers (purple). GAPDH serves as loading control.**B**) CDC42 inhibition rescues neural conversion of KLHL4-deficient iPSCs. Control or KLHL4-depleted CRISPRi iPSCs were treated and subjected to neural conversion as described above and analyzed by qRT-PCR analysis for expression of lineage markers color-coded as described above. Marker expression was normalized to control iPSCs and RPL27 was used as endogenous control. n=2 biological replicates with 3 technical replicates each. **C**) Experimental design for *in vivo* electroporations testing the ability of ^GFP^CDC42^T17N^, a constitutively inactive CDC42 variant that can bind GEFs but is deficient in recognizing effectors, to rescue KLHL4 MO phenotype. Increasing concentrations of ^GFP^CDC42^T17N^ variant were co-expressed with KLHL4 MO, while control MO with empty vector was injected on the contralateral side. Of note, while the low rescue levels of ^GFP^CDC42^T17N^ were analyzed at HH8 when the neural folds have risen, the medium and high levels were analyzed at early neural fold stage (HH7), when the control side is a flat neural plate and the neural plate border domain is less well defined, but experimental side shows risen neural folds indicating accelerated neurulation. **D)** Ectodermal patterning defect phenotype caused by loss of KlHL4 MO was rescued by co-expression of low dose CDC42 ^T17N^, but medium and high doses resulted in the appearance of accelerated neural folding as evaluated from whole mount embryos after HCR FISH by using probes for the neural plate border marker *Msx1*, the neural plate marker *Myc-N*, and the non-neural ectoderm marker *Dlx5*. Scale Bar = 250µm. Cartoon depicts a model for how an optimal amount of CDC42 signaling is required for normal ectodermal patterning and neurulation. **E**) Graph depicts ratios of anterior *Msx1* intensity of the experimental side relative and to the control side of the embryo. For reference, *Msx1* intensity rations of embryos that were injected with control MO on both sides are shown. n = 3-5 embryos per condition as indicated, error bars = s.e.m., * = p < 0.05, ** = p < 0.001, **** = p < 0.0001, one-way ANOVA)

## DISCUSSION

Differentiation processes are often driven by epigenetic, transcriptional, and translational network changes ^5, 66^. Here, we identify an essential posttranslational mechanism that regulates small GTPase signaling by converting a canonical pathway effector into an inhibitor to orchestrate key steps of early human development, a concept we hypothesize to be a common principle of regulating organogenesis. Specifically, we propose that (1) tissue-specific expression of the substrate adaptor protein KLHL4 spatiotemporally restricts monoubiquitylation activity of a CRL3 complex to the ectoderm layer and (2) monoubiquitylation by CRL3-KLHL4 balances cytoskeletal-based signaling by the ubiquitous small GTPase CDC42 and thereby coordinates ectodermal patterning and neurulation in the developing head (***Fig. 6***). Our work thus reveals an important function for ubiquitylation in coordinating cell-fate changes and morphological rearrangements during development and defines sensitive monitoring of CDC42 activity by a ubiquitin-dependent rheostat as crucial to establishing the anterior nervous system and craniofacial skeleton.

**Figure 6:**
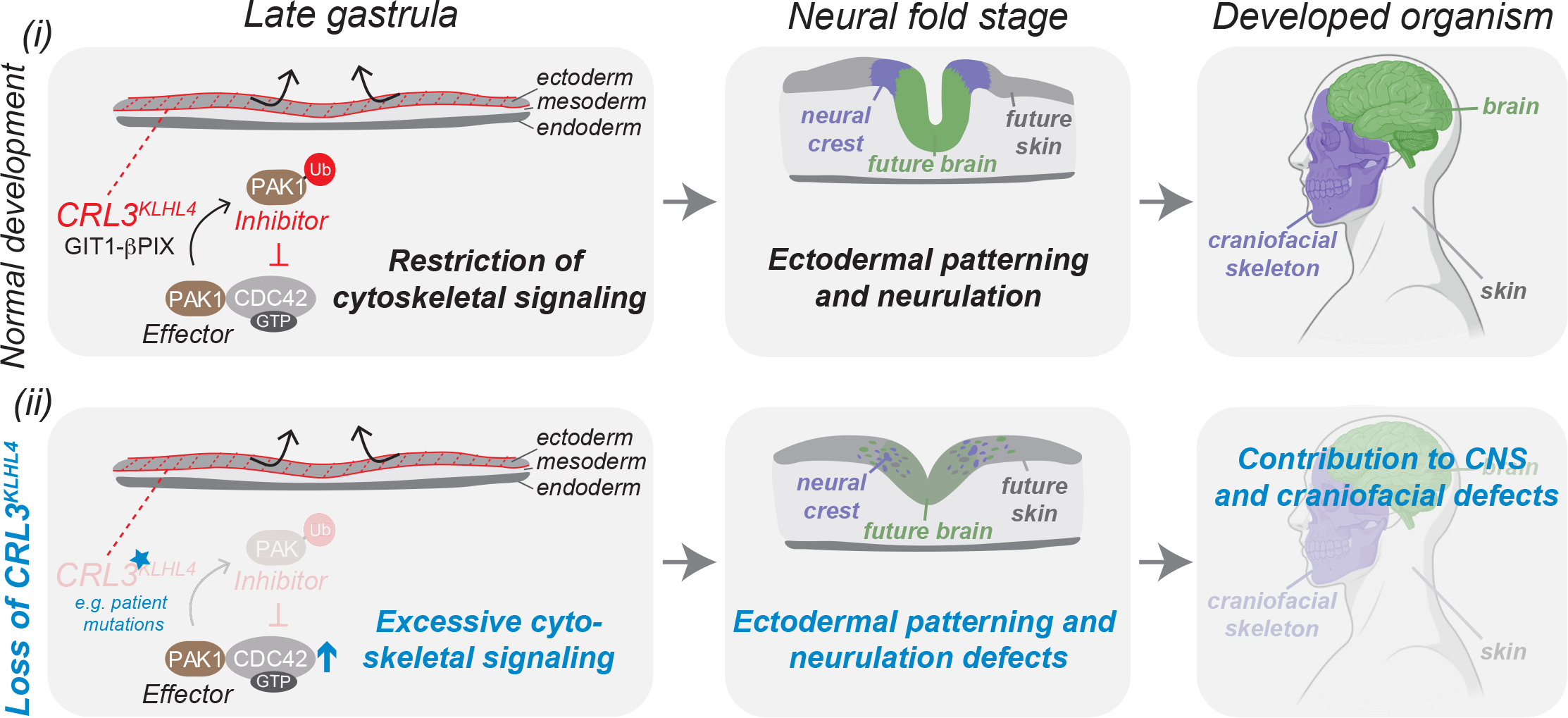
Modulation of CDC42 signaling by a ubiquitin-based effector-to-inhibitor switch coordinates ectodermal patterning and neurulation in the vertebrate head. Model depicting how CRL3-KLHL4-mediated restriction of anterior CDC42 signaling regulates ectodermal patterning. Canonically, CDC42 is activated by specific GEFs, including the GIT1-β-PIX complex, and inactivated by dedicated GAPs. The activated, GTP-bound form of CDC42 can bind and stimulate effector proteins, including PAK1, to mediate cytoskeletal and cell polarity-based downstream signaling. (i) During normal development, tissue-specific expression of CRL3-KLHL4 allows restriction of these canonical CDC42 signaling pathways in the anterior ectoderm of the gastrula. This inhibition relies on the CDC42-activating complex GIT1-β-PIX, which recruits CRL3-KLHL4 to its substrate PAK1. Ubiquitylation converts this canonical CDC42 effector kinases into a CDC42 inhibitor to restrict downstream signaling. This negative regulation ensures proper spatial ectodermal cell-fate specification, neural plate folding, and ultimately faithful development of the skin, the nervous system, and the craniofacial complex. (ii) Loss of CRL3-KLHL4 activity results in aberrant development. Hypomorphic mutations in *CUL3* or *KLHL4* impair CRL3-KLHL4 complex assembly and PAK1 ubiquitylation. This causes reduction of the negative feedback system and excessive CDC42 downstream signaling, which compromises ectodermal patterning, neural plate folding, and contributes to central nervous system and craniofacial phenotypes observed in patients with *CUL3* and *KLHL4* variants.

We show that tissue specific expression of the CRL3 substrate adaptor KLHL4 implements monoubiquitylation-based restriction of CDC42 signaling pathways to coordinate ectodermal patterning and neurulation. Intriguingly, monoubiquitylation mediated by different CRL3 substrate adaptors is also required for other steps of neural crest development during specification (via KBTBD8^35, 67^) and later during collagen secretion in cranial chondrocytes (via KLHL12^34^). We hence propose that CRL3-mediated monoubiquitylation of spatiotemporally regulated substrates is a common regulatory principle driving cell-fate determination and tissue morphogenesis that allows integration of multiple steps of the formation of the vertebrate head and possibly in the entire embryo.

Our results demonstrate that CRL3-KLHL4 is essential for brain and face formation, as neurodevelopmental and craniofacial disease-causing variants in *CUL3*^*41, 68, 69*^ reduce CRL3-KLHL4 complex formation. In addition, a novel variant in *KLHL4* (p.I287V) in a patient with severe intellectual disability, brain malformations, and craniofacial defects, was not able to support ectodermal differentiation *in vitro* or *in vivo*. Although this patient was recently diagnosed with VPS35L-associated Ritscher-Schinzel syndrome (OMIM: #618981) based on a previously unrecognized homozygous *VPS35L* variant^70^, the patient presents on the severe end of the spectrum of *VPS35L* associated phenotypes^71^ and other genetic factors might therefore contribute to the severity of the phenotype. Our model is further corroborated by previous reports that demonstrate hyperactivating variants in PAK1 and CDC42 to result in various neural and neural crest-derived deficiencies in patients, including diverse neurodevelopmental and craniofacial defects^59, 72^. Thus, an optimal amount of CDC42 activity is crucial for cranial morphogenesis. In this context, our data suggests that CDC42 restriction by CRL3-KLHL4-dependent PAK1 monoubiquitylation in the anterior ectoderm is a means to coordinate morphological rearrangements and cell-fate commitment through modulating actin cytoskeleton-associated cellular functions. Together with established roles of mechanical forces in directing embryonic stem cell behavior^73, 74^, our study therefore raises the exciting possibility that ubiquitin-dependent regulation of the actin cytoskeleton can be responsible for hard wiring cell-fates during development.

Our mechanistic analyses reveal a previously unrecognized principle of small GTPase signaling that employs a complex enzymatic logic to coordinate ectodermal patterning and neurulation. Monoubiquitylation of the canonical CDC42 effector kinase PAK1^63, 64^ results in inhibition of CDC42. Intriguingly, CRL3-KLHL4 uses the CDC42-activating complex GIT1-β-PIX as co-adaptor to recruit and monoubiquitylate PAK1, which thus directly participates in the effector-to-inhibitor conversion. Recent reports have demonstrated that GIT-PIX complexes can form molecular condensates that concentrate limited quantities of enzymes to distinct cellular compartments, including focal adhesion complexes^54,75^. Therefore, assembling activators and effectors in such membraneless compartments and utilizing monoubiquitylation to convert effectors into inhibitors may serve as a rapid means to rewire CDC42-based cytoskeletal signaling pathways in response to morphogens. Given the prevalent roles of the Rho family of small GTPases during embryogenesis^76^, we predict that similar effector-to-inhibitor conversions will be a common mechanism to balance small GTPase signaling to coordinate tissue morphogenesis and cell-fate commitment during other aspects of development.

## MATERIAL & METHODS

### Antibodies, Plasmids, Oligonucleotides

Key reagents used in this study are summarized Table S6.

### Mammalian cell culture and transfections

HEK 293T and RPE-1 cells were maintained in DMEM with 10% fetal bovine serum. Plasmid transfections were carried out using PEI. siRNA transfections were carried out with Lipofectamine RNAiMAX (Invitrogen) according to manufacturer’s instructions using 10 nM for each siRNA. Cells were routinely tested for mycoplasma using the MycoAlert Mycoplasma Detection Kit from Lonza (LT07-118).

### Pluripotent stem cell culture

iPSCs (WTC with dCas9-KRAB, AICS90) and hESCs (H1 line, WAe001-A) were maintained under feeder free conditions on Matrigel-coated plates (#354277, BD Biosciences) in mTeSR™1, (#05871/05852, StemCell Technologies Inc.). iPSCs were routinely passaged with acctuase (# 07920, StemCell Technologies Inc.), while hESCs were routinely passaged with collagenase (#07909, StemCell Technologies Inc.). Cells were routinely tested for mycoplasma using the MycoAlert Mycoplasma Detection Kit from Lonza (LT07-118).

### Lentiviral infections

Lentiviruses were produced in HEK 293T cells by cotransfection of lentiviral constructs with third generation packaging plasmids (Addgene) for 48–72 hr. Transduction was carried out by infecting 2×10^5^ hES H1 cells per well of a 6-well plate with lentiviruses in the presence of 6 μg/ ml Polybrene (Sigma) and 10 μM Y-27632 ROCK inhibitor. For transduction of lentiviruses carrying ectopic expression vectors, cells were centrifuged at 1,000g at 30C for 90min. Media was replaced with 2mL mTESR1 containing 10 μM Y-27632 ROCK inhibitor. After 4-6d of selection with appropriate antibiotic (1 μg/ml puromycin for sgRNA constructs, 200 μg/ml G418 for pINDUCER20 constructs), iPSCs or hESCs were analyzed and used in differentiation experiments.

### Neuroectodermal differentiation of iPSCs

Neural conversion of iPSCs was performed using STEMdiff™ Neural Induction Medium (#05831, StemCell Technologies Inc.) in combination with a monolayer culture method according to the manufacturer’s technical bulletin (#28044). In brief, single cell suspensions were prepared by treatment of hES cells with accutase and 1.5 – 2.0 × 10^6^ cells were seeded per well of a 6-well plate in 4mL STEMdiff™ Neural Induction Medium supplemented with 10 μM Y-27632 ROCK inhibitor. Neural induction was performed for indicated time periods with daily medium change.

### iPSC rescue experiments

iPSCs (WTC with dCas9-KRAB, AICS90) were transduced with control sgRNAs or sgRNAs targeting KLHL4 and selected and maintained in 1 μg/ml puromycin. Cells were then stably transduced with pINDUCER20 plasmids (with indicated KLHL4^FLAG^ constructs containing wobble mutations that render them resistant to sgRNA recognition or indicated ^HA^PAK1 constructs). Cells were selected and maintained with 200 μg/ml G418 for 4-5d. For the rescue experiments, these cell lines were then treated with or without 1 μg/ml doxycycline to induce construct expression and subjected to neural conversion for indicated time periods. Cells were harvested for small GTPase pull down assays, immunoblotting, RNA extraction, or fixed for immunofluorescence analysis.

### Embryoid body differentiation assays

Embryoid body (EB) formation from hESCs (H1 line) were performed using Aggrewell™800 plates (#27865, StemCell Technologies Inc.) and APEL2 medium (#05270, StemCell Technologies Inc.) following the guidelines of the manufacturer’s technical manual (#29146). In brief, single cell suspensions were prepared by treatment of hES cells with accutase (#07920, StemCell Technologies Inc.) and 1 × 10^6^ cells were seeded per well of an Aggrewell™800 plate in APEL2 medium supplemented with 10 μM Y-27632 ROCK inhibitor (#72307, StemCell Technologies Inc.) followed by 72h incubation with daily media change. Embyroid bodies were then embedded into Matrigel and differentiated in APEL2 medium for 6d. Medium was replaced every day.

### Immunoprecipitations

For co-immunoprecipitation experiments, HEK 293T or RPE-1 cells were transiently transfected with indicated FLAG-tagged, MYC-tagged, and/or HA-tagged constructs and incubated for 48 hours at 37**º**C with 5% CO_2_. Cells were harvested by scraping in 1xPBS and centrifuged at 300xg for 5 minutes. The cell pellets were either stored at -80**º**C or directly used for immunoprecipitation experiment. For each condition, typically 1×10 cm dishes (HEK293T) or 1×15cm dishes (RPE-1) were used. Cells were lysed in two pellet volumes of ice-cold lysis buffer (20 mM HEPES pH 7.3 containing 110 mM potassium acetate, 2 mM magnesium acetate, 50mM NaCl, 1 mM EGTA, 0.1% NP-40, 1x protease inhibitors (Roche), 1x Phos-Stop (Roche), and 2 mM phenanthroline. Cells were sonicated and the lysates were cleared by centrifugation at 20,000xg for 25min. To remove residual lipids, the supernatant was filtered through 0.22 um filter (Millex-GV). Subsequently, the lysates were quantified using Pierce 660nm reagent (Thermo, #22660) and equal amounts of lysates were incubated with ANTI-FLAG-M2 agarose (Sigma) for 2h at 4 °C. Beads were then washed three times with lysis buffer and eluted in 2x urea sample buffer (150mM Tris pH 6.5, 6M urea, 6% SDS, 25% glycerol and a few grains of bromophenol blue) followed by immunoblot analysis for interaction partners. For compPASS-based mass spectrometry, HEK293T cells transiently transfected with different KLHL4^FLAG^ constructs were lysed and subjected to anti-FLAG immunoprecipitation as described above (4 x15 cm dishes per condition). Beads were then washed three times with lysis buffer and eluted in lysis buffer supplemented with 0.5mg/mL 3xFLAG peptide (Sigma). Eluted proteins were precipitated by adding 20% TCA followed by overnight incubation on ice. Protein pellets were washed three times with ice-cold 90% acetone in 0.01 M HCl, air dried, and further processed for mass spectrometry analysis as described below. For immunoprecipitation of endogenous KLHL4 complexes, lysates of 2 × 15 cm dishes of hESC (H1 line) were prepared as described above. After incubation with anti-KLHL4 or control antibodies (mIgGs, Santa Cruz) at 4°C for 1h, Protein G beads (Roche) were added for 2 h. After washing with lysis buffer, bound proteins were eluted with 2x SDS sample buffer and analyzed by immunoblotting. For immunoprecipitation of ^FLAG^CUL3 complexes from hESC undergoing early stages of ectodermal differentiation, hESCs (H1 line) stably transduced with ^FLAG^CUL3 or indicated congenital disease-causing variants were subjected to neural conversion for 1d and harvested (2×15cm dishes per condition). Cell pellets were lysed in two pellet volumes of lysis buffer (20 mM HEPES pH 7.3, 150 mM NaCl, 110 mM KOAc, 2 mM Mg(OAc)_2_, 5 mM EDTA, 5 mM EGTA, 0.2% NP-40) supplemented with 2mM phenantroline and protease inhibitors (Roche) on ice followed by brief low-amplitude sonication and subsequent trituration through 25 gauge needles. Lysates were cleared by centrifugation, passaged through a 0.45 μm membrane filter, incubated with Protein G agarose for 30 mins at 4°C to remove nonspecific interactors, and incubated with ANTI-FLAG-M2 agarose (Sigma) for 1 h at 4 °C. After washing with lysis buffer, FLAG-tagged protein complexes were eluted with lysis buffer containing 0.5 mg/ mL 3xFLAG peptide in three 15 min incubations at 30°C, 800rpm. Eluted proteins were precipitated by adding 20% TCA followed by overnight incubation on ice. Protein pellets were washed three times with ice-cold 90% acetone in 0.01 M HCl, air dried, and further processed for mass spectrometry analysis as described below.

### Mass spectrometry analysis

Eluates from FLAG IPs were precipitated with TCA overnight, reduced, alkylated, separated from FLAG peptide via S-TrapTM mini columns (Protifi) and in-column digested with trypsin overnight. Tryptic digests were analyzed using an orbitrap Fusion Lumos tribrid mass spectrometer interfaced to an UltiMate3000 RSLC nano HPLC system (Thermo Scientific) using data dependent acquisition. Initial protein identification was carried out using Proteome Discoverer (V2.4) software (Thermo Scientific). To compare the impact of disease-causing variants on known CUL3 interactors, the average intensity of the top 3 peptides of each interactor was normalized against that of CUL3 for each condition. Normalized known CUL3 interactors were compared to WT CUL3 (set to 1) and plotted as a heatmap. For each CUL3 IP condition, 2 biological replicates were performed and averaged. To determine high confidence interaction partners for KLHL4, search results from Proteome Discoverer of KLHL4^FLAG^ IPs were exported into Scaffold4 and compared with approximately 30 reference immunoprecipitations against different FLAG-tagged bait proteins using a python script programmed according to the CompPASS software suite^49^.. For determination of the KLHL4 interaction network, three independent KLHL4^FLAG^ IPs were compared as replicates against the reference IPs. Thresholds for high confidence interaction partners (HCIPs) were top 5% of interactors with highest Z-score and highest WD score. To narrow down putative substrates of KLHL4 in the interaction map, we compared relative total spectral counts for each HCIP found in WT KLHL4 IPs to the ones found in KLHL4 I287V, KLHL4ΔCUL3, KLHL4^AP1-4D/E-R^, and WT KLHL4 Ips from GIT1-depleted cells (average of 2 technical replicates for each).

### Purification of recombinant proteins

^His^MBP, ^His^MBP-GIT1^WT, His^MBP-GIT1^RK3D^, GST, GST-RBD and GST-PBD were purified from Rosetta II (DE3) competent cells using previously established protocols ^76^. In brief, cells were grown at 37*°*C until an OD_600nm_ = 1.5-2.0 was obtained, chilled, and protein expression was induced with 500nM IPTG at 16*°*C for 16h. Bacteria were harvested, resuspended in lysis buffer (0.1M Tris [pH 8.0], 0.5M NaCl, 5mM EDTA, 0.1% Triton-X100) supplemented with a protease inhibitor cocktail (Roche), and lyzed in a chilled LM10 microfluidizer at 15,000 psi. Lysates were clarified by centrifugation at 4*°*C, 50,000xg for 30min and incubated with Ni-NTA agarose (Qiagen) or glutathione Sepharose (GE Healthcare) at 4C for 2h. For ^HisMBP^GIT1^WT^ and ^HisMBP^GIT1^RK3D^, beads were washed with 10-15 bead volumes of wash buffer (0.1M Tris [pH 8.0], 0.5M NaCl, 20mM Imidazole), and subsequently eluted with 5-10 bead volumes of elution buffer (0.1M Tris [pH 8.0], 0.5M NaCl, 300mM Imidazole). For GST-tagged proteins, beads were washed with 10-15 bead volumes of wash buffer (0.1M Tris [pH 8.0], 0.5M NaCl), and subsequently eluted with 5-10 bead volumes elution buffer (0.1M Tris [pH 8.0], 0.5M NaCl, 20mM Glutathione). Protein eluates were dialyzed overnight at 4*°*C in dialysis buffer (0.1M Tris [pH 8.0], 0.3M NaCl) and concentrated using amicon ultra concentrators, MWCO 30kDa (Millipore sigma) and further purified via gel filtration proteins using a Superdex 200 Increase (GL 10/300) column. Proteins were concentrated, aliquoted and snap frozen in liquid N_2_ to be stored at -80*°*C.

KLHL4^FLAG^ was purified from HEK 293T cells (10 × 15-cm dishes). Lysates were prepared and subjected to anti-FLAG immunoprecipitation as described above. To remove endogenous interaction partners, beads were washed three times with lysis buffer containing 1 M NaCl, three times with lysis buffer containing 2% NP-40, and three times with lysis buffer, followed by elution from the beads with lysis buffer containing 3xFLAG peptide (0.5 mg/ml). KLHL4 protein was quantified against BSA using commassie-stained SDS page gels.

### In vitro reconstitution of GIT1-KLHL4 binding

To reconstitute GIT1-KLHL4 binding *in vitro*, 1.2μM ^His^MBP, ^His^MBP-GIT1^WT^, and ^His^MBP-GIT1^RK3D^ were immobilized on amylose beads TB buffer (20mM HEPES [pH 7.3], 100mM KC_2_H_3_O_2_, 2mM Mg(C_2_H_3_O_2_)_2_ 1mM EGTA, 0.1% NP-40) for 30min at 4*°*C. Excess/unbound GIT1 was removed by washing once with TB buffer. 0.6μM KLHL4^FLAG^ were added in total binding volume of 200μL an incubated for 1h at 4*°*C, followed by three washes with TB buffer, elution in 2x urea sample buffer, and immunoblot analysis.

### Small GTPase pull down assays

For determining the amount of GTP-bound small GTPases, iPSCs subjected to neural conversion for 3d (1×10cm dises per condition) were washed with PBS, followed by lysis in 250μL lysis buffer (50 mM Tris pH 7.5, 10 mM MgCl2, 0.5 M NaCl, and 2% Igepal) on ice. Lysates were sonicated, cleared by centrifugation at 4C at full speed in a table top centrifuge for 10 min, quantified via Pierce 660nm protein assay. Equalized lysates were incubated with GST, GST-RBD, and GST-PBD immobilized on glutathione agarose for 1h at 4*°*C (20μL of beads at a 2mg/mL concentration). Beads were washed once with wash buffer (25 mM Tris pH 7.5, 30 mM MgCl2, 40 mM NaCl) and taken up in 2xurea sample buffer, followed by immunoblot analysis. Relative small GTPase activity was calculated by dividing the GTP-bound state present in the GST pull down fraction to the total levels of each small GTPase detected in the input fractions followed by normalization to control cells.

### Quantitative real-time PCR (qRT-PCR) analysis

For qRT–PCR analysis, total RNA was extracted and purified from cells using the NucleoSpin RNA kit (#740955, Macherey Nagel) and transcribed into cDNA using the SuperScript™ IV First-Strand Synthesis System (#18091050, ThermoFisher Scientific). Gene expression was quantified by PowerUp SYBR Green qPCR (#A25741, ThermoFisher Scientific) on a CFX96 Real-Time System (Bio-Rad). Nonspecific signals caused by primer dimers were excluded by dissociation curve analysis and use of non-template controls. Loaded cDNA was normalized using RPL27 as an endogenous control. Gene-specific primers for qRT–PCR were designed by using NCBI Primer-Blast. Primer sequences can be found in Table S5.

### Cluster analysis

mRNA abundance was measured by RT-qPCR for different conditions. The datasets were plotted as a heatmap in Python using the Seaborn library. Hierarchical clustering of samples was performed using the Bray-Curtis method with average linkage.

### Immunofluorescence microscopy

For immunofluorescence analysis, self-renewing or differentiated iPSCs or hESCs were seeded on Matrigel-coated coverslips using accutase, fixed with 4% formaldehyde in PBS for 20 min, permeabilized with 0.5% Triton in PBS for 10 min, blocked in 2% BSA for 1h and stained with indicated primary and secondary antibodies and/or Hoechst 33342 for 1 hour. Images were taken using a Nikon A1R*+* HD confocal microscope system (Nikon Instruments, Melville NY). 488 nm, 561 nm and 640 nm laser lines provided illumination for hoechst, AF 488, Rhodamine Red X and AF647 fluorophores, respectively. Data were acquired using Galvano mode at 1024×1024 with no line averaging A Z-piezo stage (Physik Instrumente USA, Auburn MA) allowed for rapid imaging in Z every 1 µm over an 8-µm Z distance. NIS-Elements (Nikon, Melville, NY) controlled all equipment. All images were maximum intensity projections and processed using ImageJ/ FIJI.

### Microinjections to chick embryos

All experiments involving animals were approved by the NIDCR Institutional Animal Care and Use Committee. The chick embryo electroporations were performed as previously described^77^. Briefly, Hamburger Hamilton (HH) stage 4 (gastrula) embryos were collected from fertilized chicken eggs on punched Whatman filter papers and placed in an electrode chamber filled with Ringer’s Solution for double sided electroporations. A glass needle pipette was filled with either the translation blocking FITC (fluorescein isothiocyanate)-conjugated Morpholino or Control Morpholino at a concentration of 1.5mM with 1 μg/μL carrier DNA, or alternatively, with an experimental plasmid with the respective empty vector for the control side. The regents were marked with two different food colors to make sure the injected area didn’t cross over the midline of the embryo. After the double injection, a complementary electrode was gently placed above the embryo in Ringer’s to apply current to deliver the material into the ectodermal germ layer. Embryos were then removed and placed into a dish with albumin to grow at 38°C and collected at HH7-8. The embryos were then checked for fluorescence and fixed in 4%PFA-PBS-0.2% Tween for 1.5h at room temperature or overnight at 4*°*C, and dehydrated by using a gradient to bring them to methanol where they were stored in -80 °C.

### In situ hybridizations

Fluorescent in situ hybridization by using Hybridization Chain Reaction (FISH-HCR). The FISH-HCR was performed as previously described ^78^ with the following modifications:

1. After applying the probes overnight, the embryos were washed for 3 hours, changing the wash buffer each hour.
2. After applying the hairpins overnight, the embryos were washed for 2 hours, changing 5xSSCT each hour, and DAPI was applied for 30 min before the final 30 min wash.
3. After the hairpins were washed, embryos were post-fixed with 4% PFA for 15 min at room temperature and washed in 0.2% PBST 2×10 min. The embryos were then imaged as wholemounts.

### Chromogenic in situ Hybridization

Whole-mount *in situ* hybridization was performed as described. Briefly, the KLHL4 probe was made by cloning the respective gene (bp 1887-2672, XM_420250.8) to a DNA vector from RT-PCR products made by using chicken whole embryo cDNA as template. The digoxigenin-conjugated RNA probes were visualized by using anti dig-AP antibody and NCB/BCIP and postfixed with 4% FA for 1H RT.

### Whole mount Immunohistochemistry on chick embryos

Immunostainings were performed as previously described ^46^. The embryos were washed with PBS and fixed with 4% PFA for overnight at 4*°*C t room and washed in 0.2% PBST 3×20 min and blocked with 5% donkey / 5% goat serum 0.2% PBST for 1h RT. Pax7 primary antibody was diluted into the blocking buffer (1/10) and nutated in slow motion for 2days at 4*°*C, washed 5x 30 min RT and incubated with the Alexa secondary antibody (1/1000) overnight at 4*°*C, washed 5x 30 min RT and mounted for imaging by using the slow-fade mounting medium (Invitrogen ProLong™ Gold Antifade Mountant).

### Cryosectioning

For cryosections, the embryos were rehydrated to 0.2%-PBST and incubated through a sucrose gradient (5% 15min RT, 15% 2-3 hr RT before incubation in 7.5% gelatin at 37*°*C for 5-7 h) followed by embedding in cryomolds. Solidified gelatin blocks were flash frozen in liquid nitrogen and stored at -80*°*C until sectioned into 20μm sections.

### IHC imaging

Imaging of the whole embryo and sections was carried out using an Andor Dragonfly 200 spinning disk confocal system coupled to a Zeiss AxioObserver (Zeiss, Thornwood, NY). A 20X apochromat air (NA 0.80) and 40X LWD apochromat water (NA 1.15) objectives were used for whole mount and cross-sectional imaging, respectively. The samples were mounted in Prolong Grid antifade mounting medium. Andor integrated laser engine provided the excitation light using 405 nm (100 mW), 488 nm (150 mW), 561 nm (100 mW), 594 nm (100 mW), 640 nm (140 mW), and 730 nm (30 mW) laser lines, and using suitable emission wavelengths for each fluorophore. A Photometrics Prime 95B CMOS (Photometrics, AZ) camera was used in 12-bit mode with 3X gain. Illumination times and relative laser intensity was varied based on sample/fluorophore brightness. Exposure times and relative laser intensity was varied based on sample/fluorophore brightness. A Z piezo stage (ASI Imaging, Eugene, OR) allowed rapid imaging in Z. Images were collected every 10 µm over 220 µm distance or every 1 µm over 40 µm distance for whole mount and cross-sections, respectively. All components were controlled by Micro-manager version 1.4.22 and was programed by ADD. Tiled images were stitched together using the Grid/Collection stitching plugin ^79^.

### Live cell imaging

The day prior to imaging RPE-1 cells expressing inducible EGFP P-tractin or Td tomato P-tractin were plated as single cells or to confluence for wound assays and treated with doxycycline (dox). The following day, cells were treated with imaging media containing Fluorobrite DMEM, 10% FBS, 100 U/ml pen/strep, 2 mM L-glutamine, 1:100 ratio of Oxyfluor and 10 mM DL-lactate to reduce photobleaching and phototoxicity. In some cases 1uM SpyDNA 650 was added to the imaging media and washout after 1 hour to visualize cell nuclei. Single cell imaging was carried out with a Yokogawa CSU-X1 confocal scanner attached to an automated Nikon Ti2 inverted microscope (Nikon, Melville, NY) using either a 60X CFI SR Plan Apo 60x water objective (NA 1.27) or a CFI Plan apo Lambda S 100XC silicone oil objective (NA 1.35). A Lun-X laser launch provided excitation for 405 nm (100 mW), 445 nm (70 mW), 488 nm (150 mW), 514 nm (150 mW), 561 nm (100 mW), and 640 nm (140 mW) laser lines, using suitable emission wavelengths for each fluorophore. A Photometric Prime 95B CMOS camera was used in 12-bit mode with 3X gain. A Piezo stage (PI) stage was used to rapidly capture Z-stacks every 0.5 µmover a 5–8-µm Z-distance. An environmental chamber surrounding the microscope maintained cells at a constant 37°C, with 10% CO2 and approximately 50% humidity (Okolabs, Tokyo). For determination of retrograde flow images were acquired every 10 seconds. Maximum intensity projections were generated and were used for actin flow quantification. For all wound assays a CFI Plan apo Lambda S 25XC silicone objective was used.

Alternatively, for wound assays a Nikon A1R HD MP system was used (Nikon Instruments). 488 nm (0.5-1.5%), 561 nm (1-2%) and 640 nm (3-5%) laser lines provided illumination for EGFP-P-tractin, td-tomato-P-tractin and SpyDNA650, respectively. Data were acquired using resonant mode and bidirectional scanning at 1024×1024 with 2X line averaging (frame rate= 7 frames/sec), and image tiling in either 3×3 or 4×3 arrays. A Z-piezo stage (Queensgate) allowed for rapid imaging in Z every 1 µm over an 8-µm Z distance. Control and knockdown cells were imaged in adjacent chambers of an Ibidi 8-well chamber (Ibidi USA, WI) and data were acquired every 15 minutes for 12-16 hours. Wounds were created using a sterile 200 um pipet tip within 20 minutes of imaging. An environmental chamber surrounding the microscope maintained cells at a constant 37°C, with 10% CO2 and approximately 50% humidity (Precision Plastics, Beltsville MD). NIS-Elements (Nikon, Melville, NY) controlled all equipment on both microscopes.

### Image Analysis and Quantification Kymograph analysis

A Fiji/ImageJ macro was created by ADD to generate three kymograph images, from which 3 slopes (distance over time) were quantified for each cell. 4D data (3D Z volumes over time) were first maximum intensity projected (MIP) prior to kymograph generation. Data were converted into microns/ sec.

### Automated cell migration tracking

For tracking of cell migration in wound assays, the SpyDNA 650 channel was MIP, then automatically contrast adjusted and converted from 12-bit to 8-bit depth. The Fiji plugin trackmate (ref PMID: 27713081) was used to track nuclei. DoG detector (object diameter: 15, quality: 2) and simple LAP tracker (settings of 15,15,2) were used for tracking, and CSV files were exported and further analyzed in Microsoft Excel, where tracks below 15 data points were not included.

### Whole mount chick embryo and anterior cross section analysis

All whole mount and sections were displayed and analyzed using maximum intensity projections of Z-stacks. Measurements were done in Fiji by taking the average fluorescence intensity in manually drawn regions of interest. Measurements of the treated side were normalized to the control side within each embryo. For length measurements of overlapping regions in sections, manual drawn regions of interest were set as ROIs in Fiji before performing a conjunction operation to find the overlapping region. Tracing of neural tube shape was performed in Fiji by manual tracing as an ROI starting from the notochord before being converted to an X-Y coordinate for display and analysis. Calculation of the neural tube fold distance was done by setting the starting/ notochord position as 0 in both X and Y coordinates before measuring the distance to the local maximum Y point.

### Exome sequencing screen for BTB variants that affect CUL3 binding

To identify candidate disease-causing variants in BTB proteins we searched exome sequencing data of patients with undiagnosed developmental diseases. We queried public databases [including DECIPHER ^80^ and denovo-db ^81^] and internal databases of the NIH and also utilized genematcher ^82^. We filtered for variants located in the BTB and BACK domain of the BTB protein. Hits were then tested for CUL3 binding by IP/IB experiments.

### Human subjects

The patient carrying the *KLHL4* variant was consented for clinical and research-based exome sequencing through the University Medical Centre Utrecht, The Netherlands.

### Quantification and statistical analysis

All experiments were repeated at least two times and statistical analyses were performed with GraphPad Prism v.9. Data are presented as mean ± SEM unless otherwise noted in figure legend. For comparisons between 2 groups an unpaired student’s t-test was applied. For comparisons between 3 or more groups, a one-way ANOVA with Tukey’s multiple comparisons test was used. Significance was set as *P* < 0.05, and denoted as: * = *P* < 0.05, ** = *P* < 0.01, *** = *P* < 0.001, **** = *P* < 0.0001. Sample sizes are indicated in the figure legends.

## ACKNOWLEDGEMENTS

We thank Vanessa Yang and Nia Teerkorpi for their technical expertise in the initial phases of this project. We thank Ken Yamada and Kylie Walters for critical feedback on the manuscript. We thank the NIDCR Imaging Core (ZIC DE000744-04) and the NIDCR Mass Spectrometry Facility (ZIA DE00075) for excellent technical assistance. This research was supported by the Intramural Research Program of the National Institutes of Dental and Craniofacial Research (NIDCR).

## AUTHOR CONTRIBUTION

Conceptualization, A.A.J., A.D., D.D.B, L.K., and A.W.; investigation, A.J.A., R.M.Y., Y.W., J.C.C., J.H., J.C., A.D.D., S.C., M.S., R.H.v.J., D.B.B, formal analysis, A.J.A., R.M.Y., Y.W., J.C.C., J.H., J.C., A.D.D., S.C., M.S., R.H.v.J., D.B.B, L.K., and A.W; supervision, Y.W., L.K., and A.W.; funding acquisition, D.B.B, L.K., and A.W; writing – original draft, A.W. and L.K.; writing – review & editing, all authors.

---

**Figure S1:**
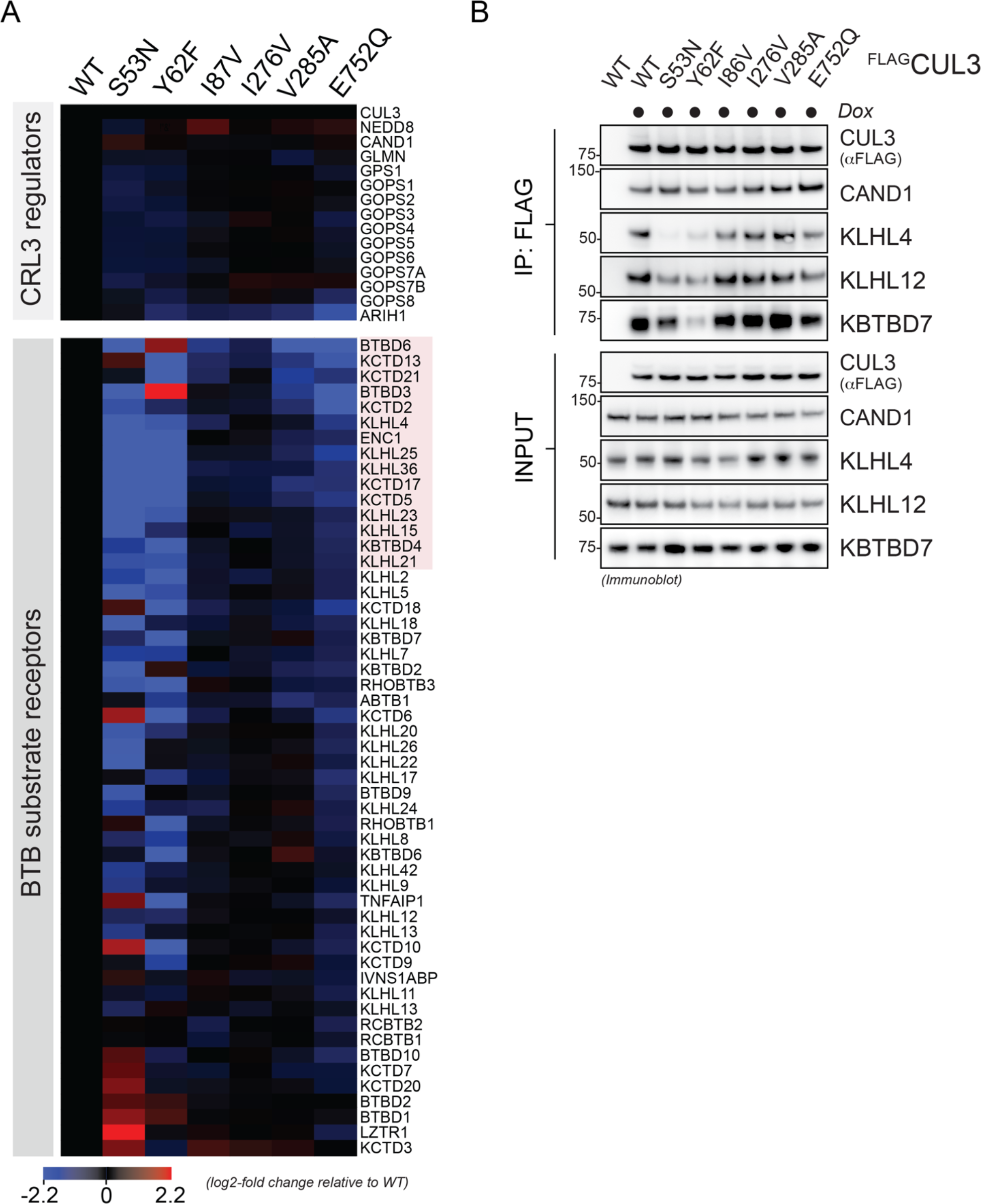
A) Neurodevelopmental and craniofacial disease-associated missense variants in CUL3 impact its interaction landscape during early stages of neuroectodermal differentiation. Neurodevelopmental and craniofacial disease-causing variants in CUL3 change interactions with BTB substrate adaptors during early stages of ectodermal differentiation. FLAG-tagged (WT) or indicated CUL3 variant were affinity-purified from hESC (H1 line) undergoing 1d of neural conversion and analyzed by mass spectrometry. Heatmap depicts the relative abundance changes in known CUL3-interacting proteins relative to WT CUL3. Highlighted in red are the 15 BTB proteins that are 1.8-fold reduced in interaction in at least 3 different CUL3 variants. n = 2 biological replicates **B**) Verification of the mass spectrometry results for a subset of CUL3 interactors. FLAG-tagged (WT) or denoted CUL3 variant were affinity-purified from hESCs (H1 line) undergoing 1d of neural conversion and analyzed by immunoblotting with the indicated antibodies.

**Figure S2:**
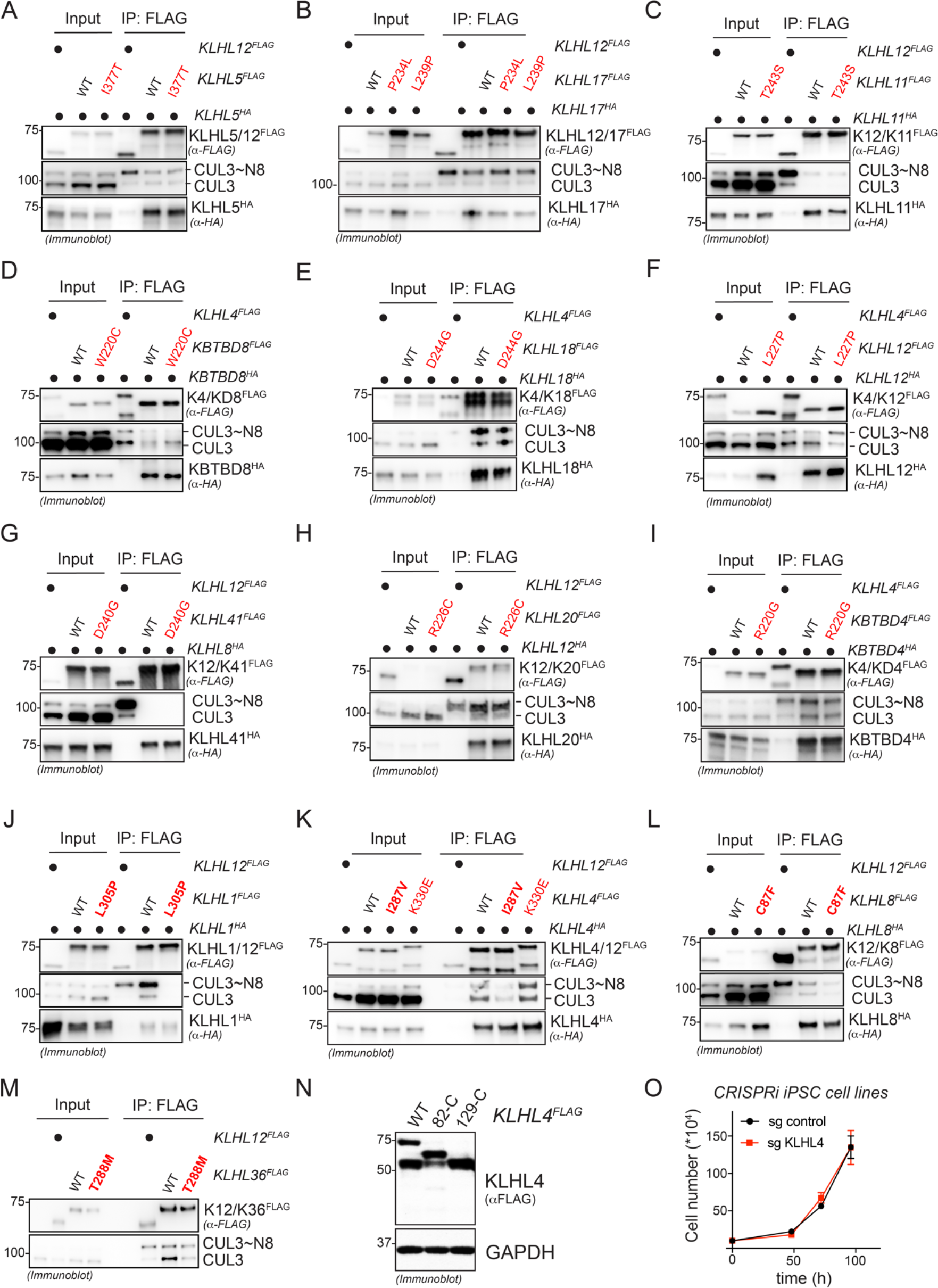
Biochemical screening of candidate disease-causing mutations from patients with undiagnosed diseases reveal several BTB variants that reduce CUL3 binding. **A**-**M**) Assessment of candidate-disease causing variants in BTB proteins identified in patients with undiagnosed diseases on their impact on BTB homodimerization and CUL3 binding. HEK293T cells were transfected with FLAG-tagged WT or indicated BTB protein variants together with HA-tagged WT BTB protein followed by anti-FLAG immunoprecipitation and immunoblot analysis with indicated antibodies. For each panel, an unrelated FLAG-tagged BTB protein was used as specificity control for dimerization. **J**) Ectopic expression of KLHL4^FLAG^ results in two isoforms in HEK293T cells. HEK293T cells were transfected with C-terminally FLAG-tagged WT KLHL4 and indicated N-terminal deletion constructs, lysed, and subjected to immunoblotting using anti-FLAG antibodies. GAPDH was used as loading control. M129 is the first methionine after M1 in the KLHL4 sequence and deletion of 128 amino acids from the N-terminus results in only the shorter isoform, while deletion of 82 amino acid still results in production of 2 isoforms. Thus, the two isoforms in WT KLHL4 likely originate from translation initiated from alternate start codons (M1 and M129). **O)** CRISPRi-mediated depletion of KLHL4 does not affect iPSC growth. Control of KLHL4-depleted iPSCs were counted at indicated times after seeding (mean of 3 biological replicates, -/+ s.d.).

**Figure S3:**
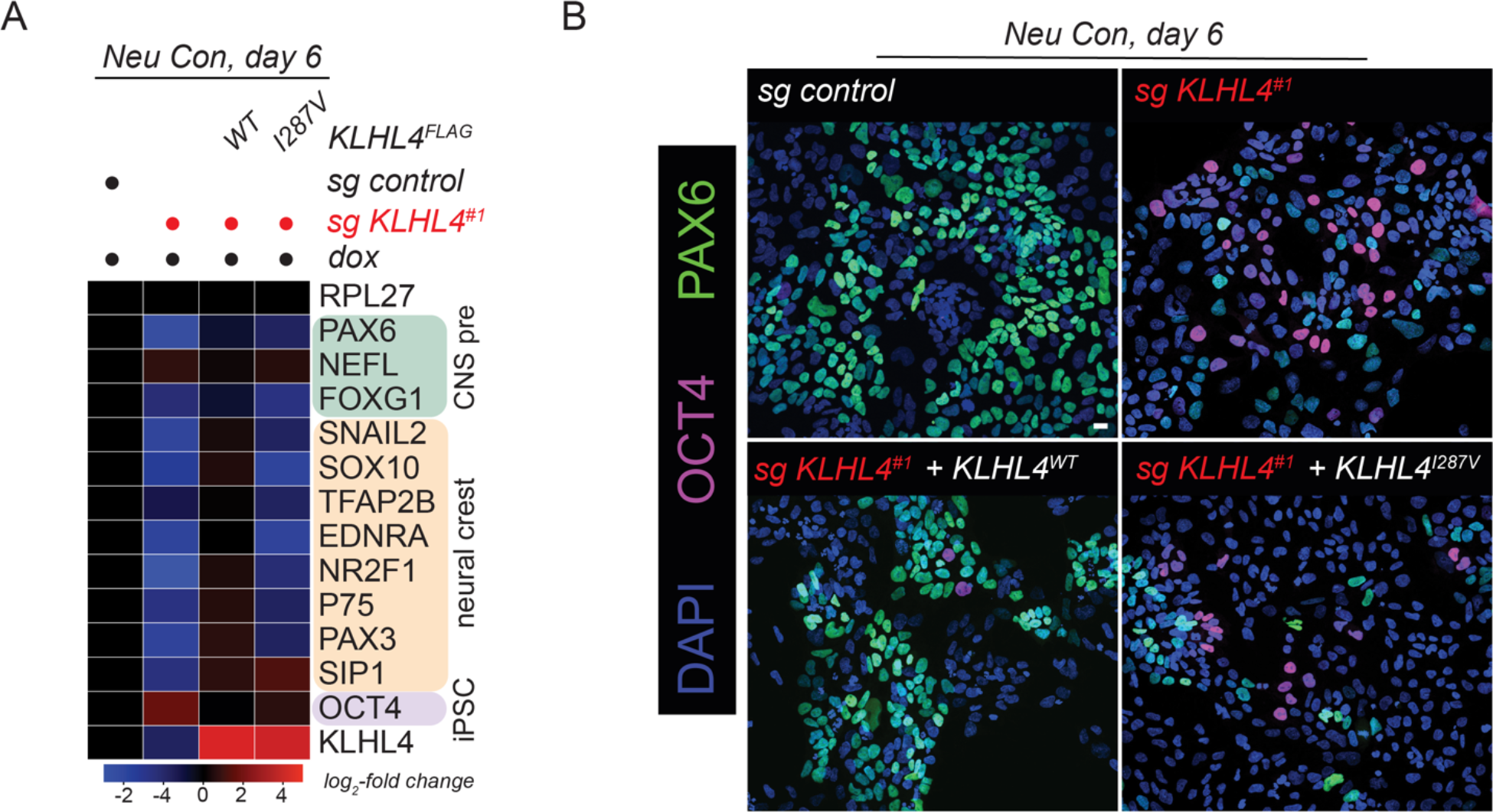
CRL3-KLHL4 ubiquitylation activity is required for neural conversion. **A**) Control or KLHL4-depleted CRISPRi iPSCs stably expressing sgRNA-resistant and doxycycline-inducible wildtype (WT) or patient variant (I287V) KLHL4^FLAG^ were generated followed by treatment with doxycycline (dox) and neural conversion for 6 days as indicated. Differentiation was monitored by qRT-PCR analysis for expression of indicated lineage markers (CNS precursors = green, neural crest = orange, iPSC = purple). Marker expression was normalized to control iPSCs. RPL27 = endogenous control. n=3 technical replicates. **H**) CRL3-KLHL4 ubiquitylation activity is required for neural conversion, as seen at the single cell level. Control or KLHL4-depleted CRISPRi iPSCs were reconstituted as described in panel A, treated with dox, and subjected to neural conversion for 8d, followed by immunofluorescence microscopy using antibodies against PAX6 (CNS precursor marker) and OCT4 (iPSC marker). Scale Bar = 20µm.

**Figure S4:**
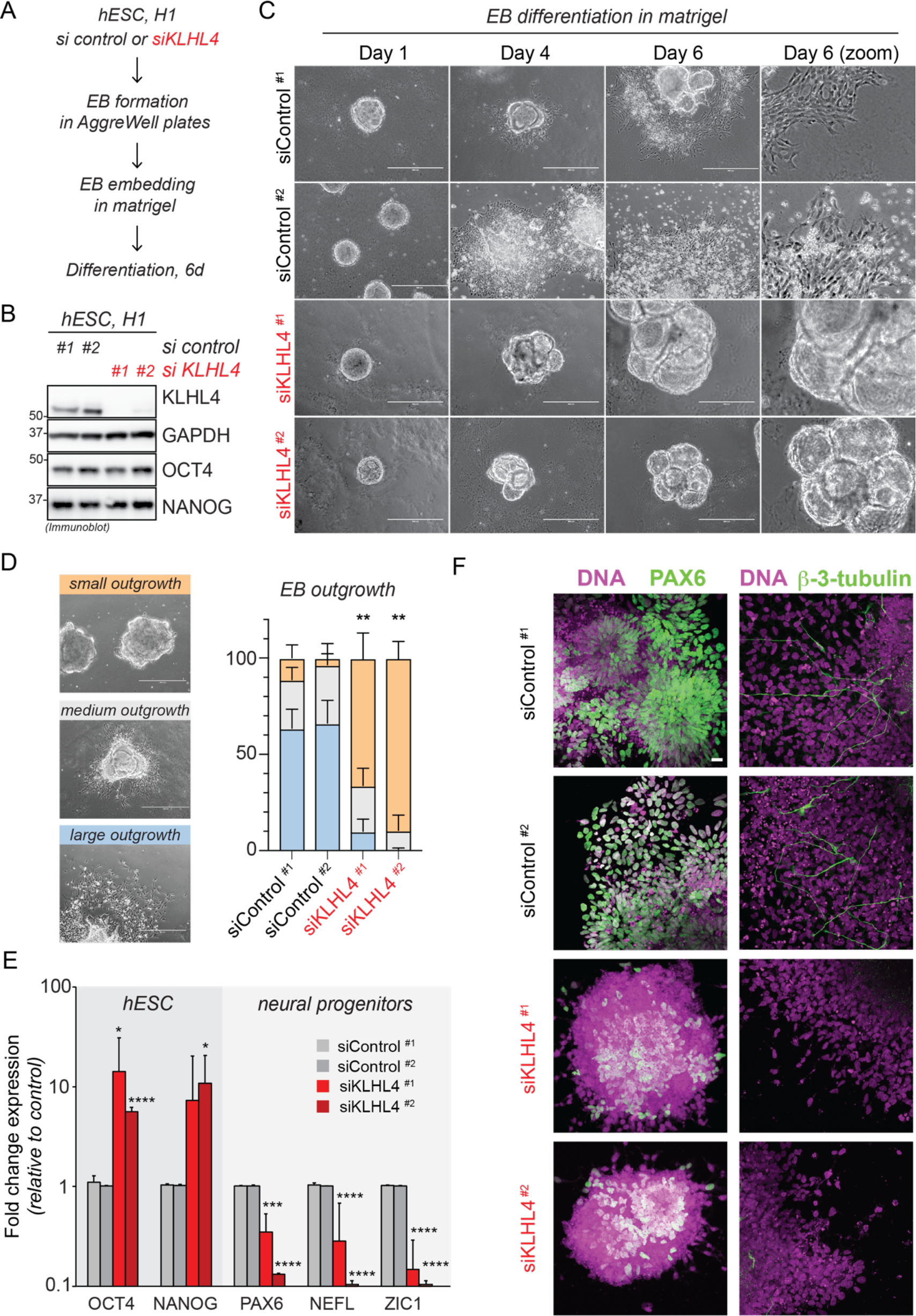
KLHL4 is required for formation of embryoid-body derived neural progenitor cells and neurons. **A**) Schematic overview of the embryoid body differentiation paradigm used to differentiate hESCs into neural progenitors and neurons. **B**) KLHL4 depletion does not alter stem cell maintenance. hESCs (H1 line) were transfected with indicated control and KLHL4 siRNA for 72h, lysed, and subjected to immunoblotting using the antibodies against the pluripotency markers OCT4 and NANOG. GAPDH = loading control. **C**) KLHL4 depletion does not interfere with embryoid body formation, but reduces cell outgrowth during differentiation. Control or KLHL4-depleted embryoid bodies were generated, differentiated as described above, and phase contrast images were taken at indicated days of differentiation. Scale bar = 400µm. **C**) KLHL4 depletion significantly reduces large cell outgrowth during embryoid body differentiation. Bar graph depicts the relative percentage of embryoid bodies with small, medium, or large outgrowth detected in each condition. Representative images of embryoid bodies with different amount of cell outgrowth are shown on the left. n= 7-11 biological replicates with 10-50 embryoid bodies counted per biological replicate, ** = p < 0.01, student t-test. **D**) KLHL4 depletion significantly reduces embryoid body-derived neural progenitor formation. Control or KLHL4-depleted embryoid bodies were differentiated for 6d as described above and subjected to qPCR analysis for the indicated hESC and neural progenitor markers. n= 3 biological replicates with 3 technical replicates each, * = p < 0.05, ** = p < 0.01, **** = p < 0.0001, student t-test. **E**) Control or KLHL4-depleted embryoid bodies were differentiated for 6d as described above and subjected to IF analysis using the neural progenitor marker PAX6 and the neuronal marker β-3-tubulin. Scale bar = 40µm.

**Figure S5:**
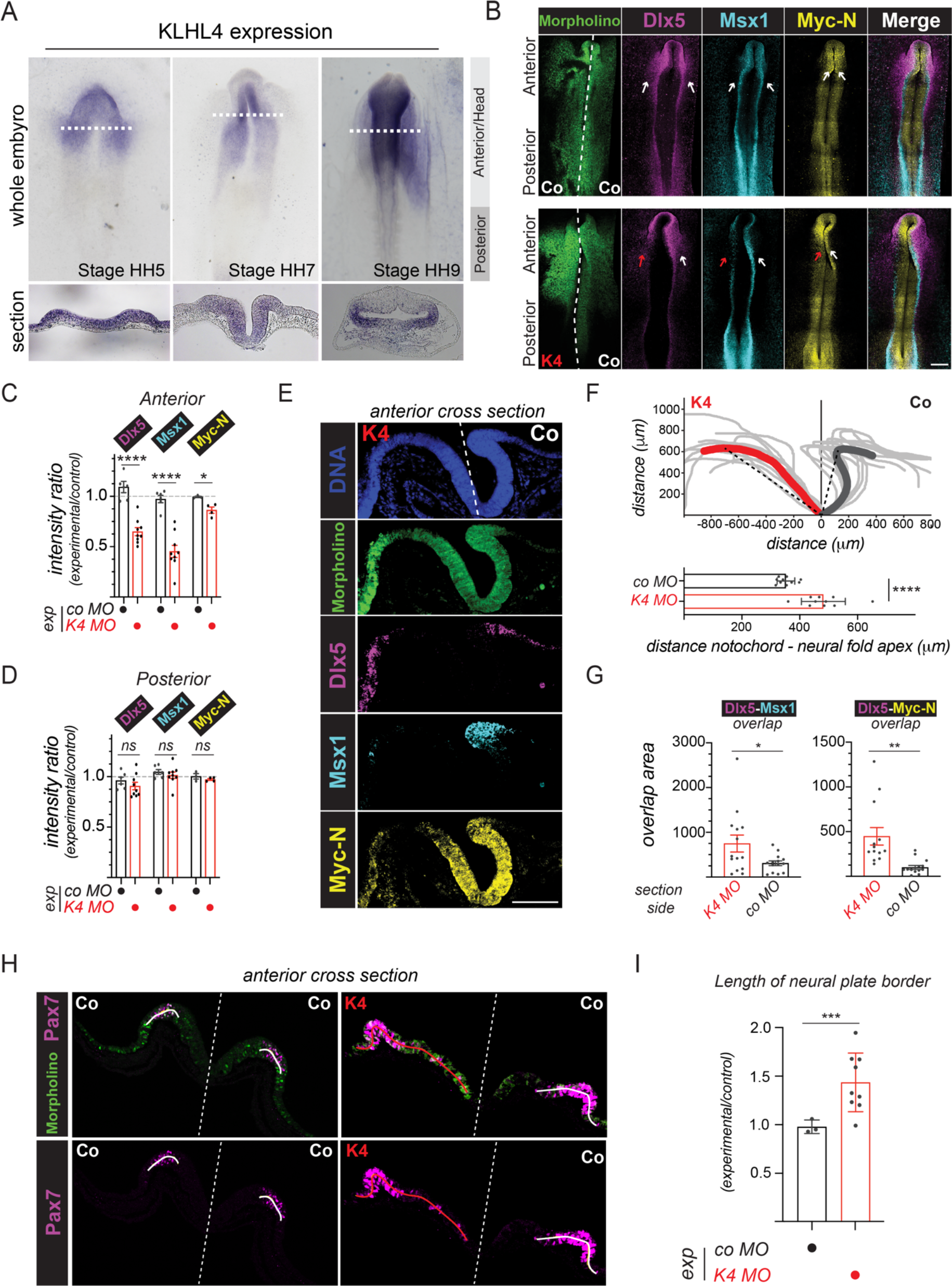
KLHL4 expression is restricted to the anterior ectoderm and CRL3-KLHL4-mediated ubiquitylation controls ectodermal patterning in the vertebrate head. **A**) Chromogenic in situ hybridization (whole embryos and corresponding cross sections indicated by dotted lines) in developing chicken embryos shows expression of KLHL4 is specific to the anterior ectoderm. After gastrulation (HH5) KLHL4 spans all domains, at neural fold stage (HH7) it is expressed in neural plate border and neural plate, and after neurulation expression is seen in the CNS portion of the neural tube. **B**) Loss of KLHL4 (K4) reduces anterior expression of markers for all ectodermal domains, including non-neural ectoderm (Dlx5), neural plate border (Msx1), and neural plate (Myc-N), whereas control embryos show no difference between sides as shown by whole mount HCR fluorescent in situ hybridization. Of note, the anterior part of the Co/Co-MO injected embryo is also shown in Figure 1G. Scale Bar = 250µm **C**) Whole mount quantification of the ratio of fluorescence intensity between experimental and control side shows significant decrease due to KLHL4 knockdown for Dlx5 (n=6 or 9 embryos as indicated, **** = p < 0.0001, unpaired t-test), Msx1 (n=6 or 9 embryos as indicated, p < 0.0001, unpaired t-test) and Myc-N (n=4 embryos per condition, p = 0.0163, unpaired t-test) at anterior axial levels. **D**) KLHL4 knockdown shows no phenotype in the trunk axial level (Dlx5: n=6 or 9 embryos as indicated, p= 0.4443, unpaired t-test; Msx1: n=6 or 9 embryos as indicated, p= 0.4691 unpaired t-test; Myc-N: n=4 embryos per condition, p= 0.2771, unpaired t-test). **E**) Cross-section from midbrain region shows reduced expression of Dlx5, Msx1, and Myc-N together with increased overlapping expression of markers between domains and a compromised folding on KLHL4 MO side as compared to control side. Scale Bar = 100µm. **F)** Tracing of neural tube shape from anterior cross-sections shows compromised folding during neurulation. Grey lines indicate individual traces and thick lines indicate average traces of control MO (dark grey) or KLHL4 MO (red) injected embryo sides. Notochord was set to 0. Bar graph shows significantly longer distance of the line from the notochord to the highest point (apex) of the neural fold on the KLHL4 MO side (n=10, p<0.0001, unpaired t-test). **G**) Measurements of overlapping length between neighboring domains show significant increase on the KLHL4 MO side as compared to control side electroporated with control MO (Dlx5-Msx1 overlap: n=14 sections, * = p < 0.05, unpaired t-test; Dlx5-MycN overlap: n=13, ** = p < 0.01, unpaired t-test). **H**) Morpholino mediated knockdown of KLHL4 increases the width of the neural plate border region as evaluated by immunostaining of *Pax7*. **I**) Measurements of the length of the domain that spans the *Pax7*-positive nuclei, indicating the neural crest region, is significantly increased on the KLHL4 knockdown side as shown by the ratio of the two sides (0.97, n=3) as compared to the ratio between two control sides (1.43, n=9, p=0.001, unpaired t-test).

**Figure S6:**
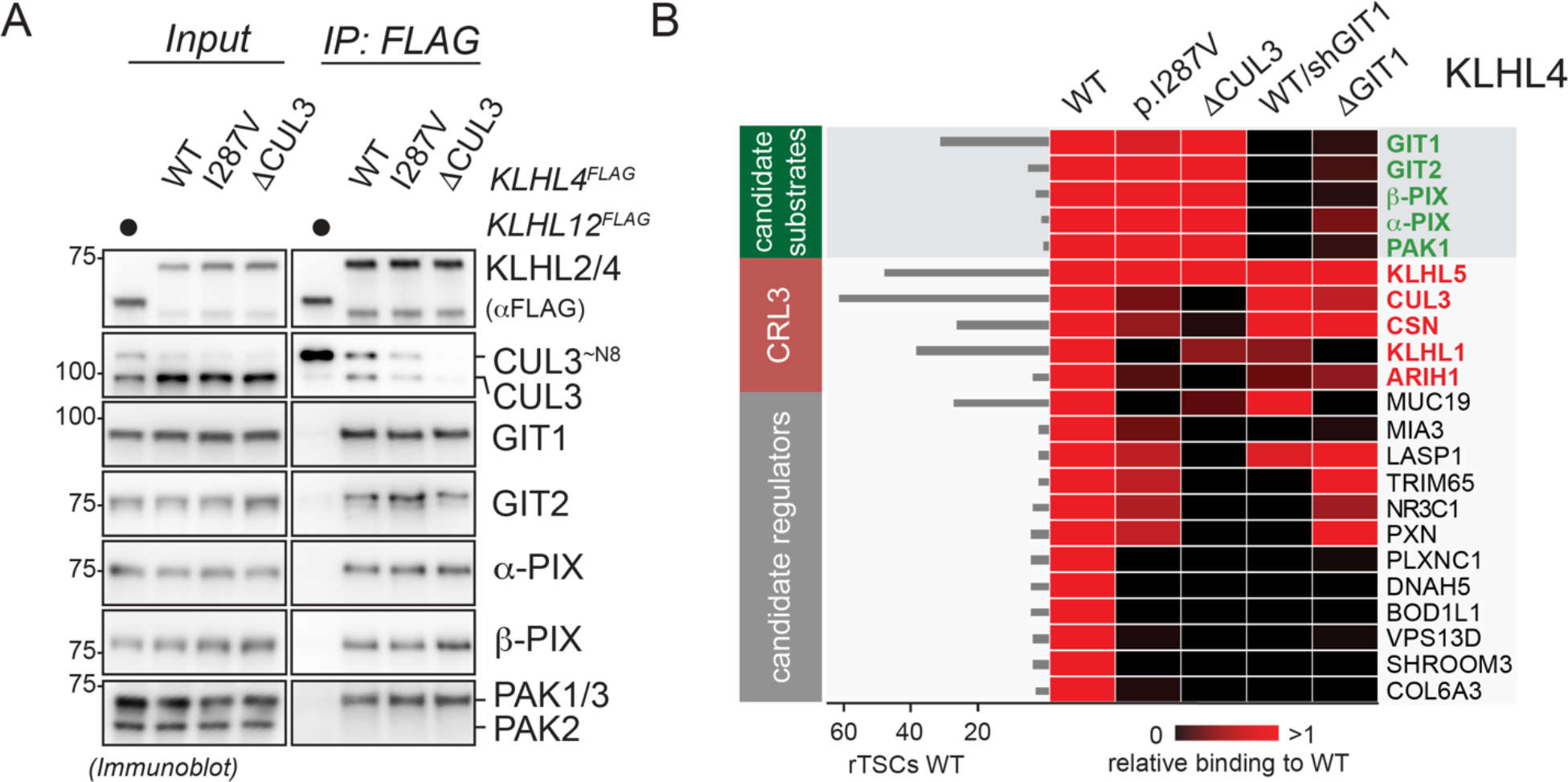
Components of the GIT-PIX-PAK signaling complex are candidate substrates of CRL3-KLHL4. **A**) Components of the GIT-PIX-PAK signaling module interact with CRL3-KLHL4 through the substrate adaptor, as shown by immunoblot analysis of indicated KLHL4^FLAG^ construct IPs from HEK293T cells. **B**) KLHL4 interacts with the GIT-PIX-PAK module through GIT1. FLAG-tagged wild-type KLHL4 (WT), patient variant KLHL4 (I287V), a severe CUL3-binding mutant of KLHL4 (ΔCUL3), and a GIT1-binding deficient mutant of KLHL4 ((ΔGIT1) were affinity-purified from HEK293T cells. In addition, FLAG-tagged wild-type KLHL4 (WT) was affinity-purified from HEK293T cells depleted of GIT1 (sh GIT1). Anti-FLAG IP fractions were analyzed for binding partners by mass spectrometry and CompPASS analysis. The heatmap depicts the relative binding of interactors identified for wild-type KLHL4 in the respective KLHL4 variant (black = no interaction, red = equal or more interaction).

**Figure S7:**
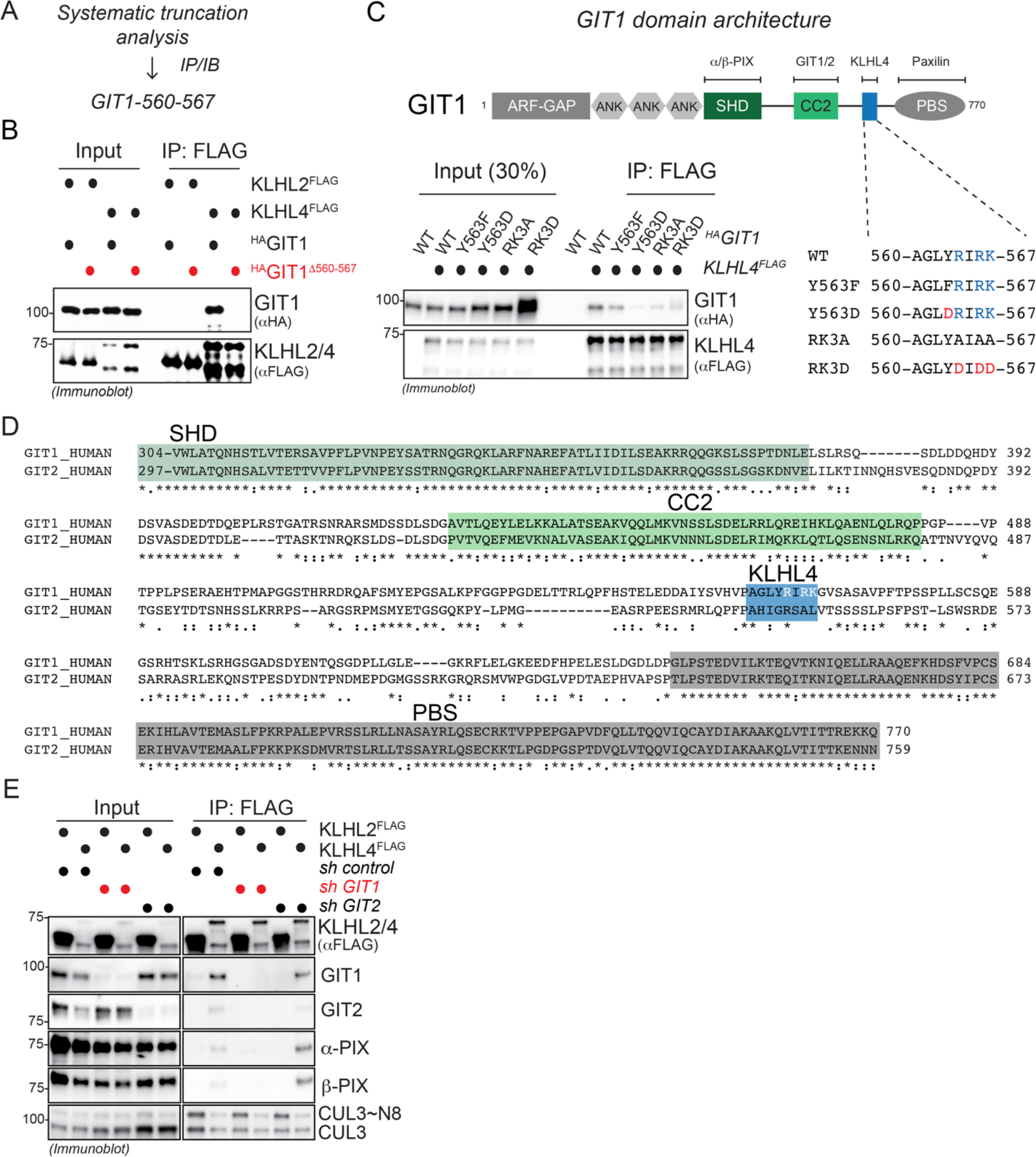
KLHL4 binds to GIT1 through a positively charged 6-amino acid stretch that is not conserved in GIT2. **A**) GIT1 truncation analysis via IP/IB experiments from HEK293T cells reveals GIT1-560-567 as region for KLHL4 interaction. **B**) Amino acids 560-567 in GIT1 are required for interaction with KLHL4. HEK293T cells were co-transfected with FLAG-tagged WT KLHL4 and HA-tagged GIT1 or GIT1Δ560-567 lysed, and subjected to anti-FLAG IP followed by immunoblotting with indicated antibodies. KLHL2^FLAG^ was used as a specificity control for interaction. **C**) Three positively charged residues in GIT1 are essential for binding to KLHL4. HEK293T cells were transfected with KLHL4^FLAG^ and different HA-tagged GIT1 constructs (WT and denoted mutants in the 8-amino acid motif that we have identified to be required for KLHL4 binding), lysed, and subjected to anti-FLAG IP followed by immunoblotting with indicated antibodies. Residue numbers in GIT1 model refer to protein isoform 3 on UniProt **D**) Amino acids 560-567 in GIT1 are not conserved in its close homologue GIT2. Sequence alignment (generated using *clustal*) showing the C-terminal part of GIT1 and GIT2 with structural domains and KLHL4 interaction site highlighted in different colors. **E**) GIT2 is not required for binding of CRL3-KLHL4 to the GIT-PIX-PAK signaling module. Control, GIT1-depleted, or GIT2-depleted HEK293T cells were transfected with KLHL4^FLAG^ or KLHL2^FLAG^, lysed, and subjected to anti-FLAG IPs followed by immunoblotting with indicated antibodies.

**Figure S8:**
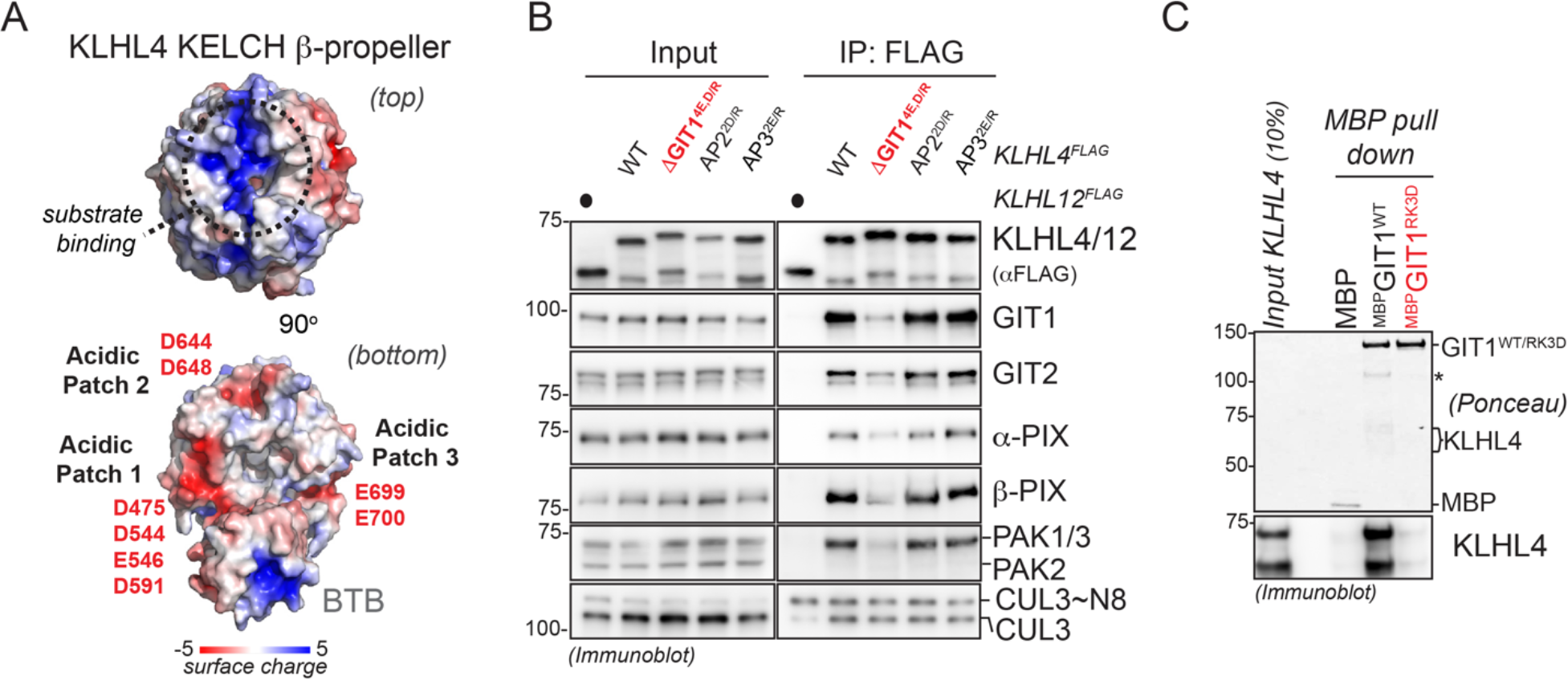
KLHL4 engages the GIT-PIX-PAK signaling module via a direct electrostatic interaction between the bottom of its KELCH β-propeller and GIT1. **A**) KLHL4 possesses three acidic patches on the bottom of its KLECH propeller. Depicted is a surface charge map of a structural model of KLHL4. The left panel shows the top of the KLECH β-propeller, the canonical binding site for substrates in other BTB-KELCH proteins, which exhibits a positively charged groove. The right panel depicts the bottom of the KELCH β-propeller that contains three acidic patches and lists the negatively charged residues contributing to them. **D**) Acidic patch 1 (denoted ΔGIT1) but not acidic patch 2 or 3 (denoted AP2 and AP3, respectively) in KLHL4 mediates binding to the GIT-PIX-PAK signaling module. HEK293T cells were transfected with FLAG-tagged KLHL12, KLHL4 (WT), and denoted KLHL4 variants with charge swap mutations in the acidic patches on the bottom of the KLECH β-propeller, lysed, and subjected to anti-FLAG IPs followed by immunoblotting with indicated antibodies. **C**) KLHL4 directly interacts with GIT1. KLHL4^FLAG^ purified from HEK293T cells under stringent conditions was incubated with recombinant MBP-tagged GIT1^WT^ or GIT1^RK3D^ and subjected to MBP pull down, followed by immunoblotting.

**Figure S9:**
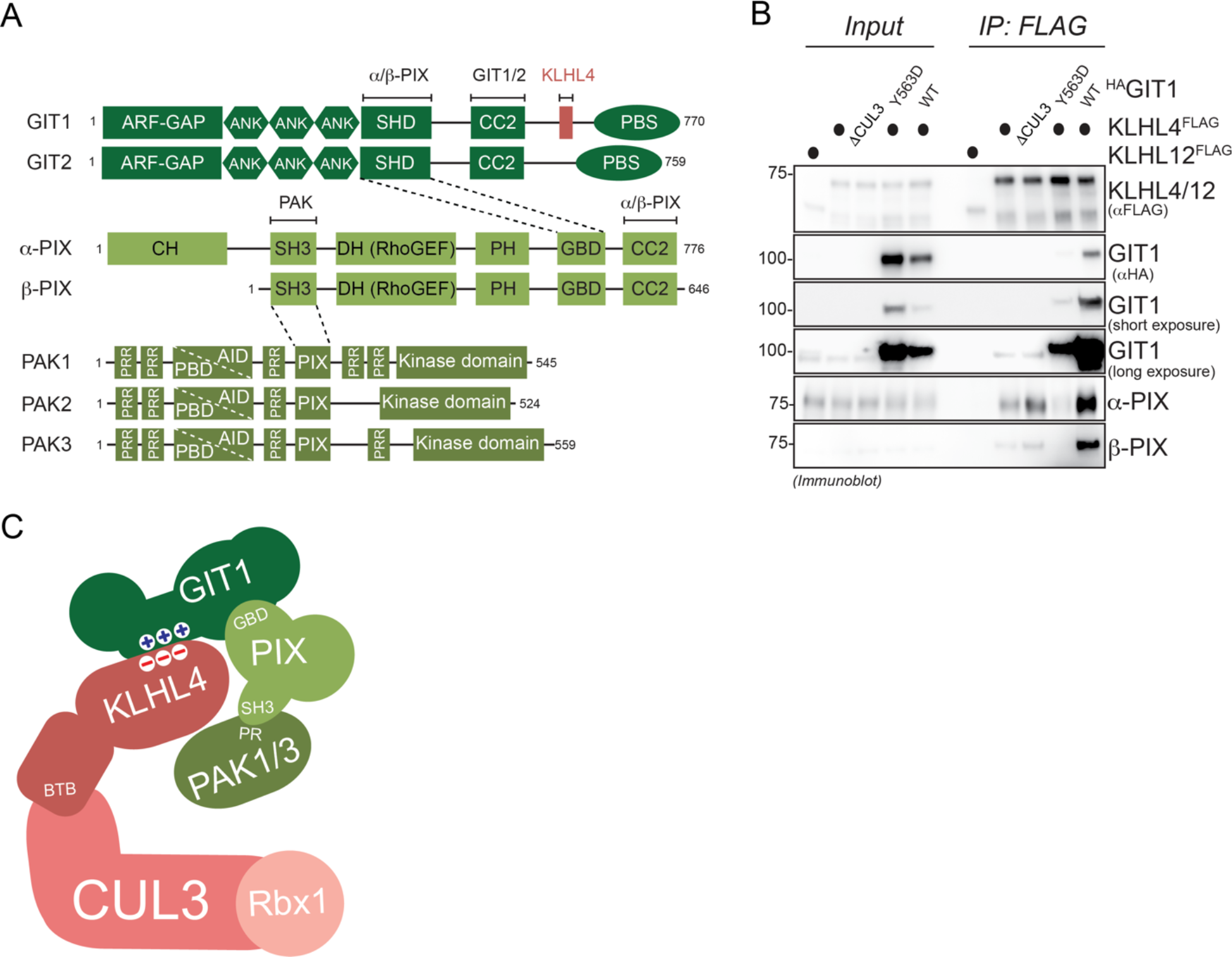
CRL3-KLHL4 utilizes GIT1-PIX complexes as substrate co-adaptors to recruit PAK1. **A**) Cartoon depicting domain structures of GIT, PIX, and group I PAK proteins. Highlighted are the known interaction domains through which GIT-PIX-PAK complexes can form. GIT proteins bind to the GIT-binding domain (GBD) of PIX proteins. GIT-PIX complexes can further associate to a proline-rich motif in group I PAKs (denoted as PIX) via SH3 domains of PIX proteins. KLHL4 can associate to these assemblies via a binding site in GIT1 (colored in red). **B**) GIT1 recruits PIX proteins to KLHL4. HEK293T cells were co-transfected with indicated FLAG-tagged KLHL4 or HA-tagged GIT1 constructs, lysed, and subjected to anti-FLAG IP followed by immunoblotting with indicated antibodies. KLHL12^FLAG^ served as specificity control for interactions. Compared to KLHL4^FLAG^ alone, co-expression of ^HA^GIT1^WT^ increases binding of PIX proteins, while co-expression of the KLHL4-binding deficient ^HA^GIT1^Y563D^ decreases binding of PIX proteins, indicating that they are sequestered. **C**) Cartoon depicting a simplified model on how CRL3-KLHL4 uses GIT-PIX complexes as co-adaptors to bind PAK1.

**Figure S10:**
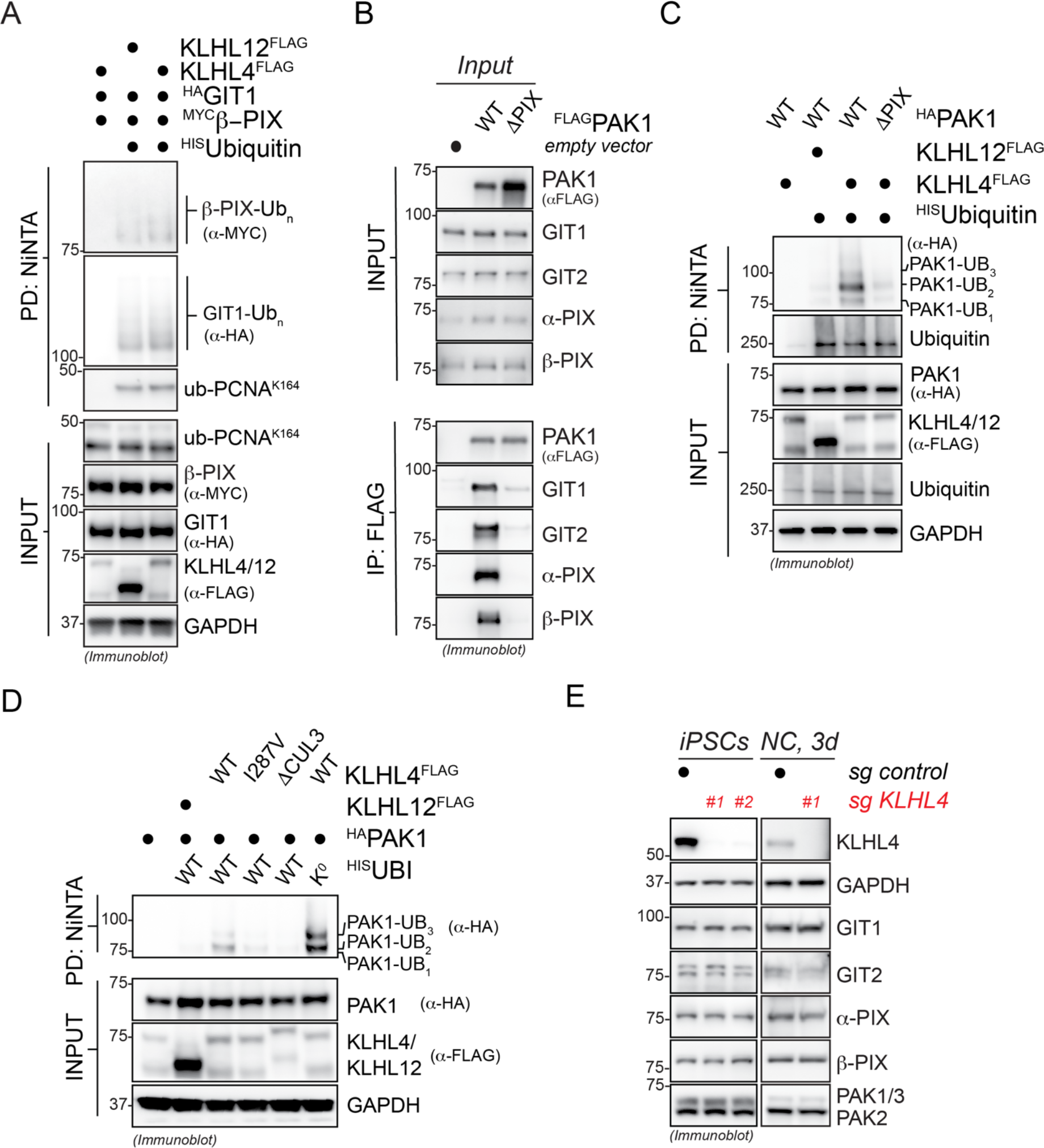
CRL3-KLHL4 utilizes GIT1-PIX complexes to recruit and multi-monoubiquitylate PAK1. **A**) GIT1/β-PIX complexes do not undergo CRL3-KLHL4-dependent ubiquitylation in cells. Ubiquitylated proteins were purified under denaturing conditions from RPE-1 cells expressing ^HIS^Ubiquitin and denoted combinations of ^HA^GIT1/^MYC^β-PIX, KLHL12^FLAG^, and KLHL4^FLAG^. Modified proteins were detected by immunoblotting using the indicated antibodies. PCNA is ubiquitylated by a distinct E3 ligase and hence is used as a control for general ubiquitylation efficiency. **B**) Verification that a previously published PIX-binding-deficient PAK1 mutant cannot engage GIT-PIX complexes in cells. RPE-1 cells were transfected with ^FLAG^PAK1^WT^ or ^FLAG^PAK1^PRP192-194AAA^ (ΔPIX), lysed, and subjected to anti-FLAG IP followed by immunoblotting with indicated antibodies. **C**) PAK1 requires interactions with GIT-PIX complexes to be ubiquitylated by CRL3-KLHL4 in cells. Ubiquitylated proteins were purified under denaturing conditions from RPE-1 cells expressing ^HIS^ubiquitin and ^HA^PAK1 (WT) or GIT-PIX-complex-binding-deficient ^HA^PAK1 (ΔPIX) in the absence or presence of KLHL4^FLAG^. Modified proteins were detected by immunoblotting using the indicated antibodies. Anti-Ubiquitin blots serve as a control for general ubiquitylation efficiency **D**) CRL3-KLHL4 multi-monoubiquitylates PAK1 in cells. Ubiquitylated proteins were purified under denaturing conditions from RPE-1 cells expressing ^HIS^Ubiquitin WT or chain formation-deficient ^HIS^Ubiquitin (K^0^) and denoted combinations of ^HA^PAK1, KLHL12^FLAG^, and KLHL4^FLAG^ variants. Modified proteins were detected by immunoblotting using the indicated antibodies. WT and chain formation-deficient ^HIS^Ubiquitin support the same pattern of PAK1 modification, suggesting that these are multi-monoubiquitylation events. **E**) KLHL4 depletion does not change steady state levels of components of the GIT-PIX-PAK signaling module. Control or KLHL4-depleted CRISPRi iPSCs or cells undergoing neural conversion for 3d were lysed and subjected to immunoblotting using the indicated antibodies.

**Figure S11:**
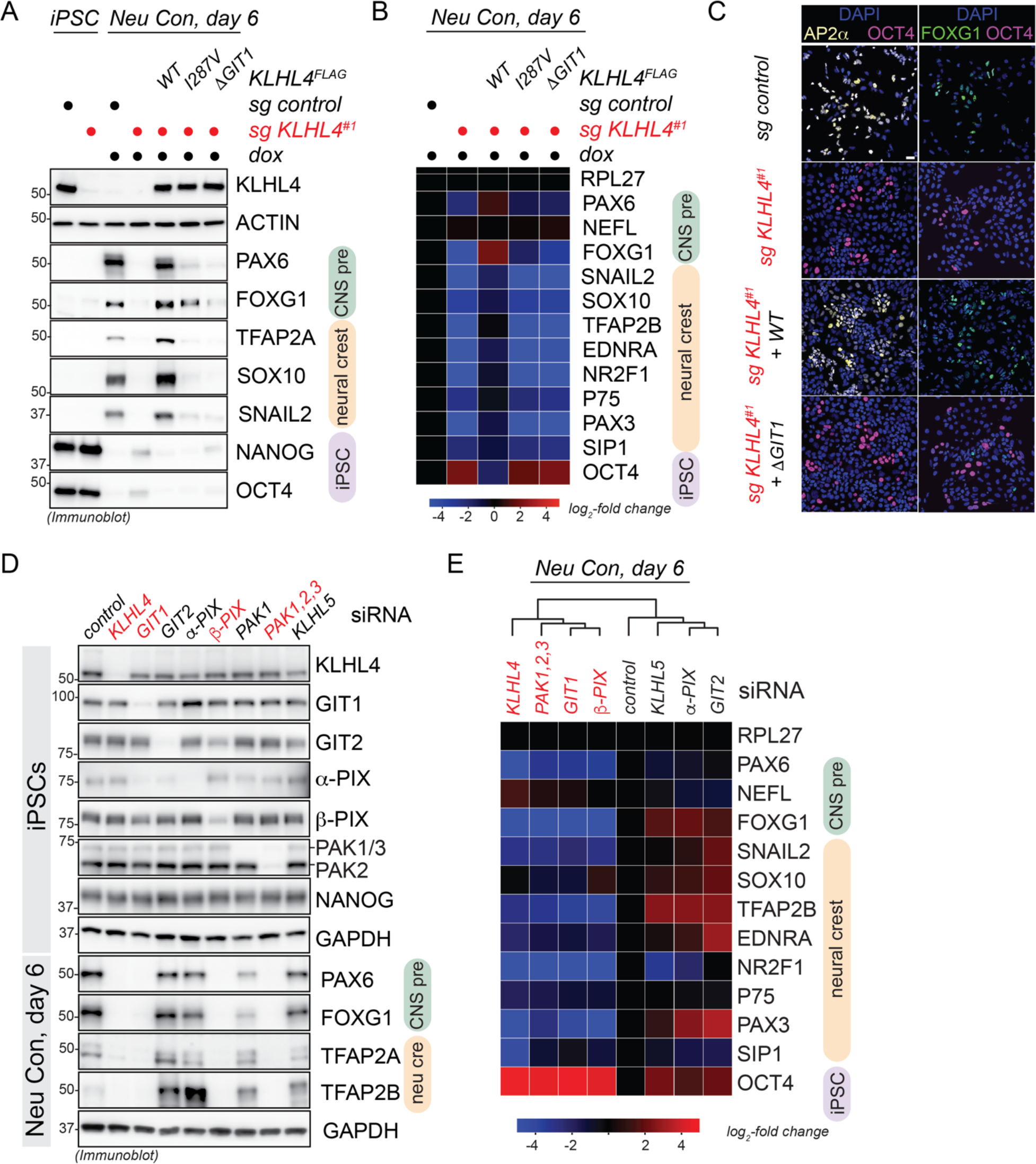
CRL3-KLHL4 controls ectodermal differentiation through the GIT-PIX-PAK signaling module. **A**) The GIT1-binding deficient mutant of KLHL4 is not able to support neural conversion. Control or KLHL4-depleted CRISPRi iPSCs stably expressing sgRNA-resistant and doxycycline-inducible wildtype KLHL4^FLAG^ (WT), patient variant KLHL4^FLAG^ (I287V), or GIT-PIX-PAK-binding deficient KLHL4^FLAG^ (ΔGIT1) were treated with doxycycline (dox) and subjected to neural conversion for 6 days. Differentiation was monitored by immunoblotting using indicated antibodies against lineage markers (iPSC = purple; CNS precursor = green; neural crest = orange; ACTIN = loading control). **B**) Control or KLHL4-depleted CRISPRi iPSCs were reconstituted and differentiated as described in panel A, followed by qRT-PCR analysis for expression of indicated lineage markers. Marker expression was normalized to control iPSCs. (RPL27 = endogenous control, n=2 biological replicates with 3 technical replicates each). **C**) Control or KLHL4-depleted CRISPRi iPSCs were reconstituted as described in panel A, treated with dox, and subjected to neural conversion for 8d followed by immunofluorescence microscopy using indicated antibodies. Scale Bar = 40µm. **D**) Depletion of GIT1, β-PIX, and group I PAKs (PAK1,2,3) phenocopies the aberrant neural conversion program observed upon KLHL4 reduction. Control CRISPR iPSCs were depleted of endogenous KLHL4 or indicated proteins using siRNA, subjected to neural conversion for 6d, and analyzed by immunoblotting. Conditions that phenocopy KLHL4 depletion are highlighted in red. **E**) Control CRISPR iPSCs were depleted of endogenous KLHL4 or indicated proteins using siRNA and cells were subjected to neural conversion for 6d, followed by qRT-PCR analysis for expression of CNS precursor markers (green), neural crest markers (orange), and iPSC markers (purple). Marker expression was normalized to control iPSCs and RPL27 was used as endogenous control. Heatmap depicts the average of 2 biological replicates with 3 technical replicates each.

**Figure S12:**
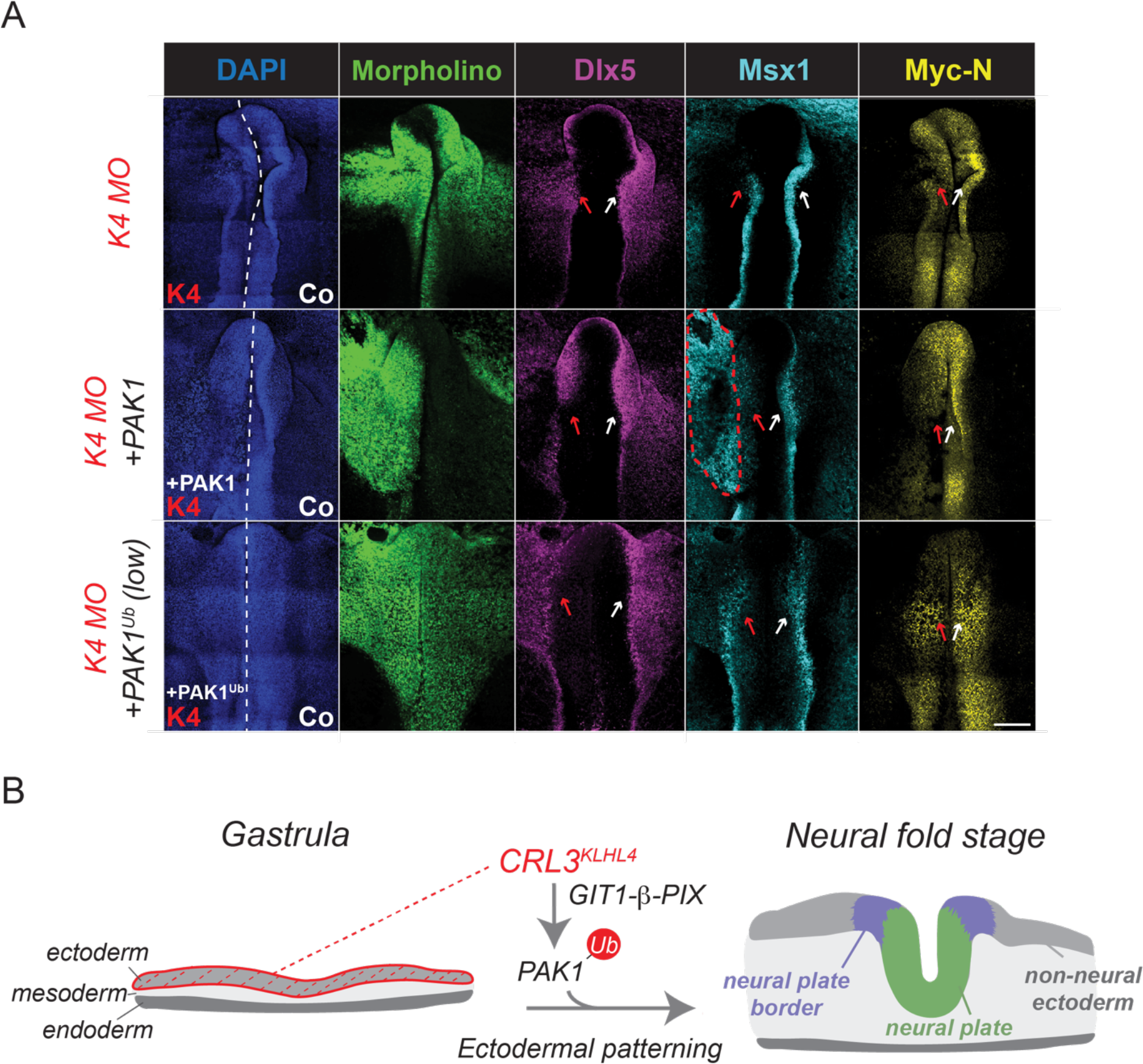
CRL3-KLHL4 controls ectodermal development through monoubiquitylation of PAK1. **A**) Co-expression of ubiquitylated PAK1 rescues the loss of KLHL4 phenotype in a dose-dependent manner as shown by expression of *Dlx5, Msx1* and *Myc-N*, the respective markers for the non-neural ectoderm, neural plate border/neural crest and neural plate as indicated by the arrows in whole mount embryos after HCR FISH. Morpholino-induced loss of KLHL4 phenotype is not rescued by the non-ubiquitylated WT PAK, as shown by decreased expression similar to KLHL4 MO alone. Additionally, expression of PAK1 induces ectopic expression of Msx1 next to the neural plate border region (circled area). Scale Bar = 250µm **B**) Model illustrating how CRL3-KLHL4, expressed specifically in the anterior ectoderm of the developing embryo, utilizes GIT1-β-PIX complexes as co-adaptors to ubiquitylate PAK1 to coordinate ectodermal domain specification and neurulation.

**Figure S13:**
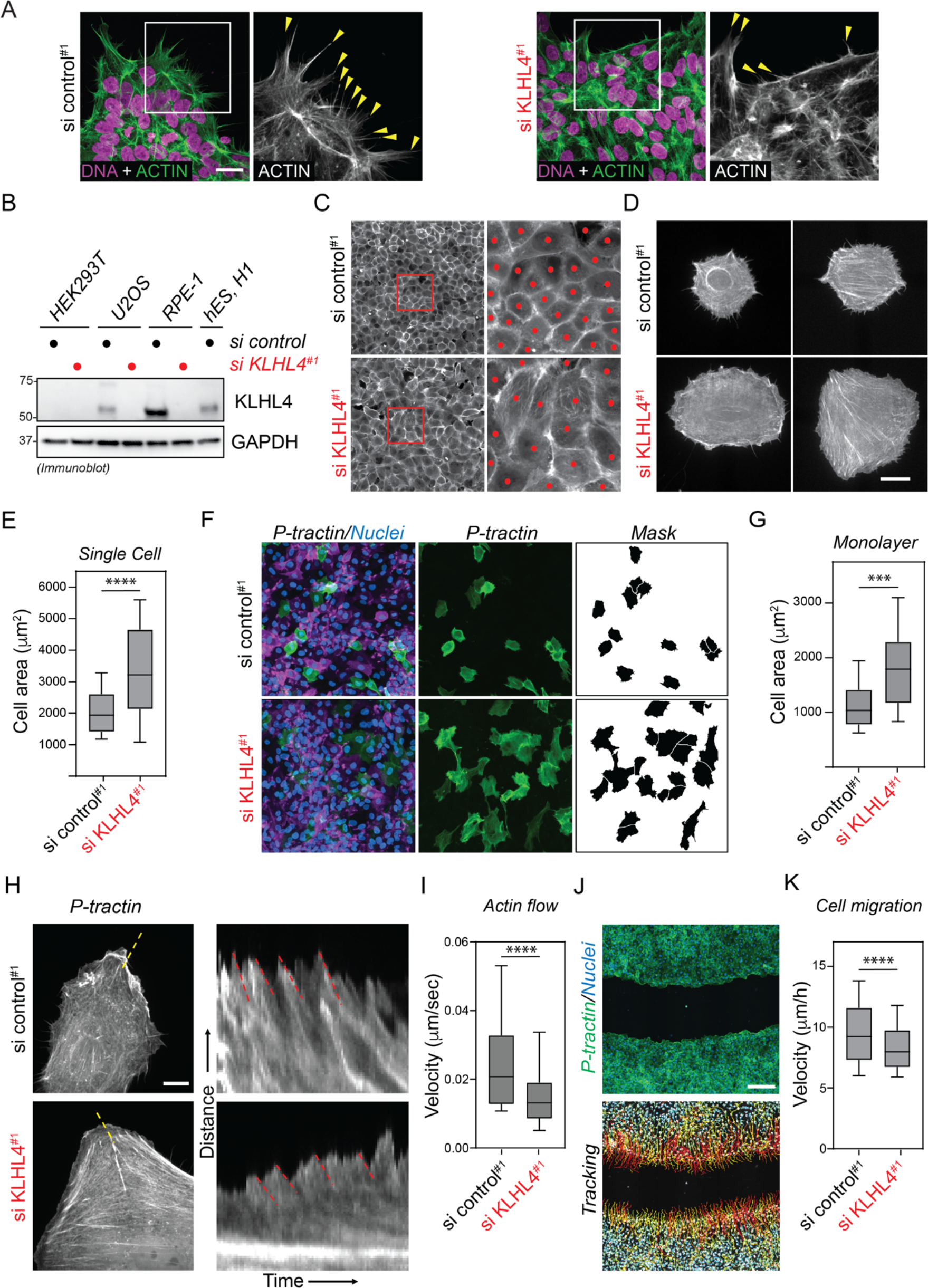
KLHL4 controls epithelial cell morphology and actin flow rates. **A**) KLHL4 regulates colony morphology and actin protrusions (arrowheads in inset), as evidence by immunofluorescence microscopy of control and KLHL4-depleted hESCs (H1 line). Scale Bar = 20µm. **B**) RPE-1 cells express high levels of KLHL4. Anti-KLHL4 immunoblot analysis of cell lysates transfected with control or KLHL4 siRNA for 72h. GAPDH = loading control. **C**) Representative examples of si control^#1^ (top) and si KLHL4^#1^ (bottom) RPE-1 cells expressing EGFP-P-tractin, demonstrating the change in cell area and concomitant cell packing common in cell monolayers. **D**) Representative examples of si control^#1^ (top) and si KLHL4^#1^ (bottom) RPE-1 cells expressing EGFP-P-tractin, demonstrate the change in cell area is also observed in single cell plating experiments. **E**) Quantification of single cell area between si control^#1^ and si KLHL4^#1^ treated RPE-1 cells. N=4, n>36. *p*≤0.0001, student t-test. **F**) Examples of si control^#1^ (top) and si KLHL4^#1^ (bottom) RPE-1 cells plated as a monolayer expressing either EGFP (green) or mScarlet (magenta)-P-tractin in a 1:10 ratio to allow for quantification of cell spread area. Nuclei (Spy 650 DNA) are shown in blue. EGFP-P-tractin cells (middle) were thresholded, masked and segmented (right) to demonstrate the enlarged cell area.**G**) Quantification of monolayer cell area between Si control^#1^ and si KLHL4^#1^ treated RPE-1 cells. N=4, n>280. *p*≤0.0001. student t-test. **H**) Si control^#1^ (top) and si KLHL4^#1^ (bottom) RPE-1 cells expressing EGFP-P-tractin were imaged at high resolution rapidly over time to observe the dynamic flow of extending lamellipodia. Dashed yellow lines mark the position each kymograph (right) was taken. Dashed red lines indicate the local flow rate of actin during protrusion/retraction cycles of the lamellipodia. **I**) Quantification of actin flow rate between si control^#1^ and si KLHL4^#1^ treated RPE-1 cells. N≥10 cells, n≥59 kymographs. *p*=0.0002, student t-test. **I**) Example image on a monolayer wound assay showing si KLHL4^#1^ treated RPE-1 cells (top) expressing EGFP-P-tractin (green) with marked nuclei (Spy650 DNA, blue). Bottom images shows the nuclear tracking used to quantify cell migration rate. **J**). Quantification of cell migration rates between si control^#1^ and si KLHL4^#1^ treated RPE-1 cells. N≥4, n≥10000 cells. *p*≤0.0001, student t-test. Scale bars: B; 20 µm, D; 100 µm, F; 10 µm, I: 200 µm.

**Figure S14:**
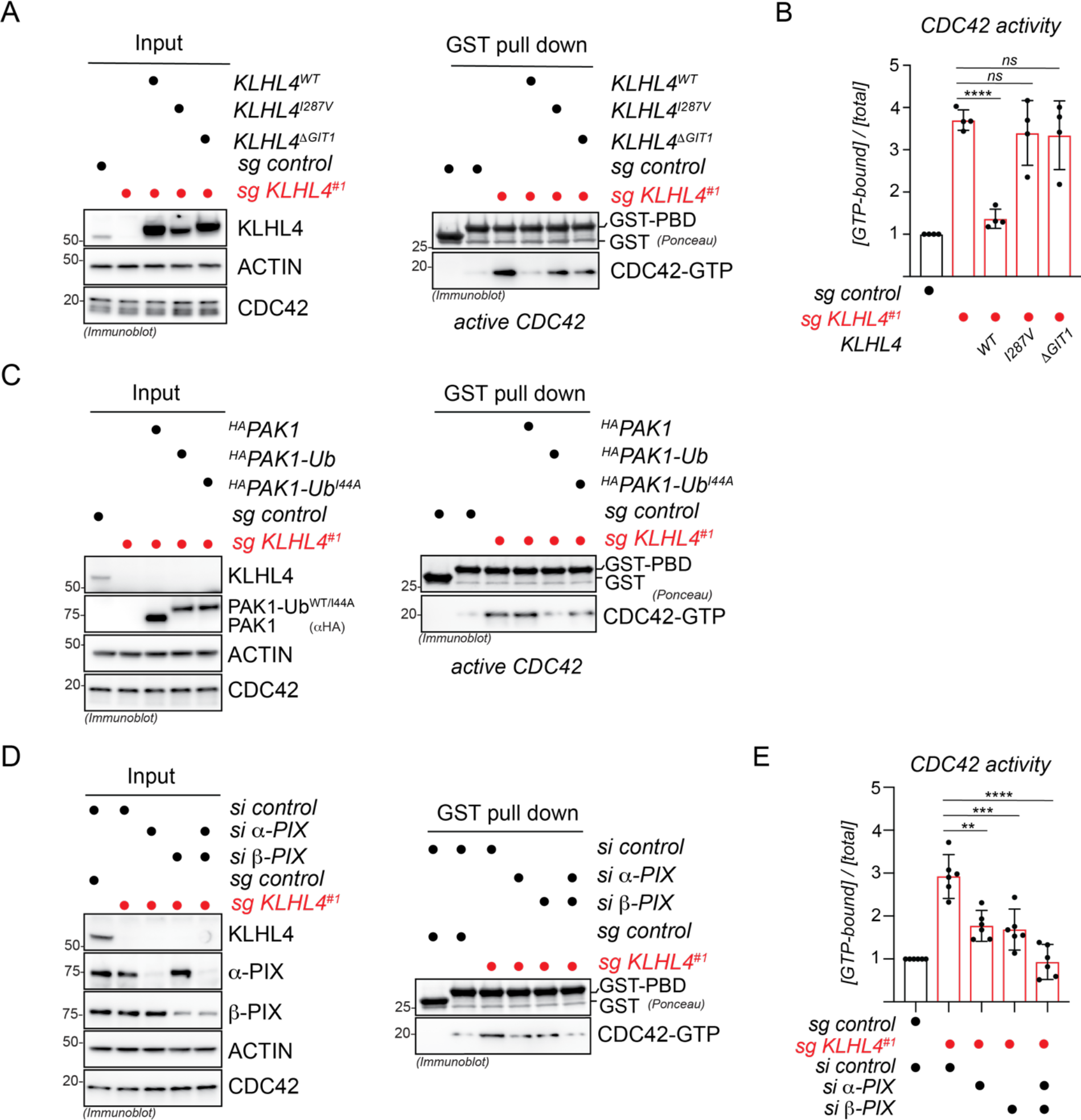
CRL3-KLHL4 mediated PAK1 ubiquitylation restricts CDC42 activity during ectodermal differentiation. **A**) CRL3-KLHL4 ubiquitylation activity and binding to the GIT-PIX-PAK module are required to restrict CDC42 activity. Control or KLHL4-depleted CRISPRi iPSCs were reconstituted with dox-inducible wildtype KLHL4^FLAG^ (WT), CUL3-binding deficient, patient variant KLHL4^FLAG^ (I287V), or GIT-PIX-PAK-binding deficient KLHL4^FLAG^ (ΔGIT1), treated with dox, and subjected to neural conversion for 3d. GTP-bound CDC42 was affinity-purified using GST-PBD followed by immunoblotting with indicated antibodies. GST was used as specificity control for binding. **B**) Quantification of the relative CDC42 activity of the experiment in panel B. n= 2 biological with 2 technical replicates each, error bars denote s.d., *** = p < 0.001, one-way ANOVA. **C**) Ubiquitylated PAK1 can restore normal levels of CDC42 activity in KLHL4-deficient cells undergoing early stages of neural conversion. Control or KLHL4-depleted CRISPRi iPSCs were reconstituted with dox-inducible ^HA^PAK1, ^HA^PAK1-Ub, or ^HA^PAK1-Ub^I44A^, treated with dox and subjected to neural conversion for 3d. GTP-bound CDC42 was affinity-purified using GST-PBD followed by immunoblotting with indicated antibodies. GST was used as specificity control for binding. Quantifications of immunoblots are shown in Figure 3E. **D**) Co-depletion of α- and β-PIX can restore normal levels of CDC42 activity in KLHL4-deficient cells undergoing early stages of neural conversion. Control or KLHL4-depleted CRISPRi iPSCs were depleted of endogenous α- and/or β-PIX using siRNAs and subjected to neural conversion for 3d. GTP-bound CDC42 was affinity-purified using GST-PBD followed by immunoblotting with indicated antibodies. GST was used as specificity control for binding. **E**) Quantification of the relative CDC42 activity of the experiment in panel E. n= 2 biological with 3 technical replicates each, error bars denote s.d., * = p < 0.01, *** = p < 0.001, **** = p < 0.001, one-way ANOVA.

**Figure S15:**
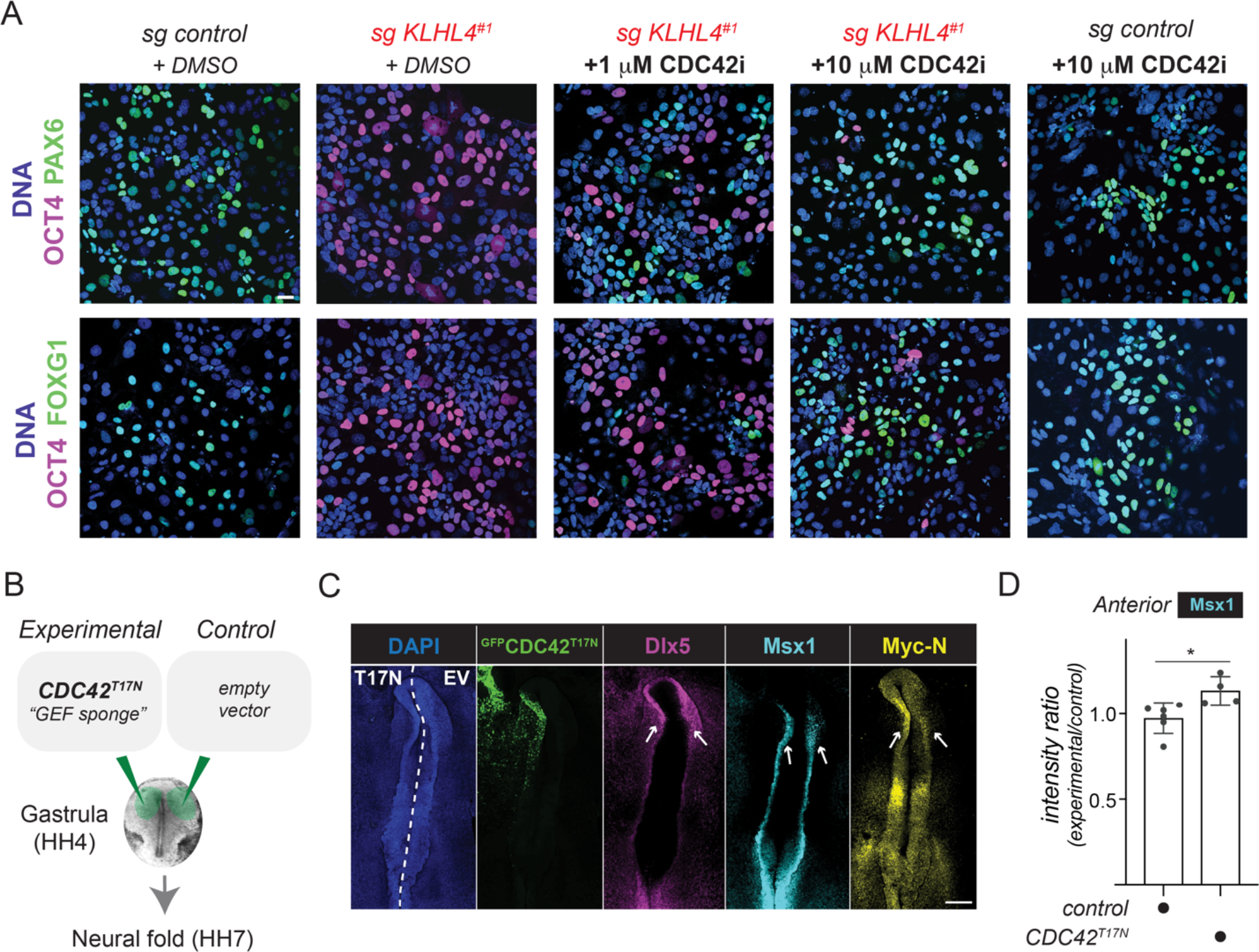
KLHL4-depletion induced ectodermal differentiation defects can be rescued by CDC42 inhibition. **A**) CDC42 inhibition rescues neural conversion of KLHL4-deficient iPSCs. Control or KLHL4-depleted CRISPRi iPSCs were treated with DMSO or indicated concentrations of the CDC42 inhibitor ML141 (CDC42i), subjected to neural conversion for 8d, and analyzed by immunofluorescence analysis for expression of the CNS precursor markers PAX6 and FOXG1 and the pluripotency marker OCT4. **B**) Expression of downstream signaling-deficient CDC42^T17N^ leads to aberrant ectodermal patterning during chick development. ^GFP^CDC42^T17N^ was electroporated into the epiblast in one side of the embryo (experimental) and an empty vector on the other side (control). **C**) Embryos injected as described in panel E were grown until the neural fold stage (HH stage 7) and analyzed by HCR using probes for the neural plate border marker *Msx1*, the neural plate marker *Myc-N*, and the non-neural ectoderm marker *Dlx5*. Scale Bar = 250µm **D**) Graph depicts ratios of anterior Msx1 intensity of the experimental side relative and to the control side of the embryo. n = 4-6 embryos per condition as indicated, error bars = s.e.m., * = p < 0.05, unpaired t-test.

