## Supplementary material for "A tissue-specific ubiquitin switch coordinates brain, craniofacial, and skin development": Note S1

**Supplementary Note 1**

Subject 1 is a male child born of non-consanguineous caucasian parents. Pregnancy and delivery were uncomplicated and was born at 40+2 weeks with a birth weight of 2665 grams (7%,, SD, -1.5), height 49 cm (32%,, SD -.47), and head circumference 35 cm (66%, SD .42). Apgar scores after 5 and 10 minutes were 7 and 9. He had distinctive features with a relative large skull with wide sutures and fontanels, a small face with pointed nose, hypertelorism, micrognathia, low-set ears, hypermobile joints, irregular inplant of his toes, a single umbilical artery, and a small penis. He also has a perimembraneous ventricular septal defect. He has partial ocular albinism (no retinal pigment, little pigment in the iris) and bilateral congenital glaucoma, resulting in blindness besides some light perception. His psychomotor development was severely delayed. He has no speech, and understands some phrases.

He had failure-to-thrive, resulting in a severe short stature with dystrophic build. He had very frequent ENT and pulmonary infections, and has hearing loss after recurrent ear infections. At the age of 9 years, he had meningococcal meningitis. He was diagnosed with a common variable immunodeficiency. At the age of 9 years, head circumference was 52.3 cm (-0.4 SD). At 19 years, his height was 137 cm (-6.5 SD), his weight 28 kg (BMI 14.9 (-4,4 SD)).

CT cerebrum showed hypoplasia of the cerebellum and vermis, a megacysterna magna, a cavum septum pellucidum, and cerebral migration defects. He developed immune thrombopenic purpura (ITP) at the age of 17 years, followed by severe autoimmune hemolytic anemia six years later.

Karyotype was 46,XY, SNP array analysis didn't show CNVs. Sanger sequencing of *CASP10, FAS*, and *FASLG* for CVID/ALPS. Trio-WES revealed a maternal missense variant in *KLHL4* (NM_019117.4(KLHL4):c.859A>G p.(Ile287Val) ChrX(GRCh37):g.86873066A>G) and a homozygous variant in *VPS35L* (c.2627C>T: p.P876L).

He has one healthy brother. His other brother was affected and had a similar phenotype as he has, with a severe psychomotor retardation, a large VSD, recurrent (mainly pulmonary and skin) infections, a relative large skull with wide sutures and a small face with low-set ears, some hypertelorism, and retrognathia. He was dysmature at birth, with an average height and head circumference. His head circumference increased rapidly due to a severe communicating hydrocephalus. He had a growth retardation. Ophthalmologic examination was normal. At the age of 21 months, he died of a sepsis after cardiac operation. CT cerebrum showed some cerebellar hypoplasia. He had a normal male karyotype and no additional genetic testing was performed or possible to date.
