## Supplementary material for "A tissue-specific ubiquitin switch coordinates brain, craniofacial, and skin development": Table S6

### Table S6: Key resources

| REAGENT or RESOURCE | SOURCE | IDENTIFIER |
|---|---|---|
| Antibodies | | |
| Mouse monoclonal anti-beta-ACTIN (clone C4) | MP Biomedicals | Cat#:691001; RRID: AB_2335127 |
| Rabbit monoclonal anti-HA-Tag (C29F4) | Cell Signaling | Cat#:3724; RRID: AB_10693385 |
| Mouse anti-HA.11 antibody | BioLegend | Cat#:MMS-101P |
| Rabbit monoclonal anti-Flag DYKDDDDK Tag | Cell Signaling | Cat#:S2368 |
| Mouse monoclonal anti-Flag DYKDDDDK Tag | Sigma | Cat#:F3165 |
| Mouse monoclonal anti-FLAG clone M2 | Sigma-Aldrich | Cat#:F1804; RRID:AB_262044 |
| Mouse monoclonal anti-MYC | Cell Signaling | Cat#:2040; RRID: AB_2148465 |
| Rabbit polyclonal anti-CUL3 | Bethyl | Cat#:A301-109A; RRID:AB_873023 |
| Rabbit monoclonal anti-GAPDH | Cell Signaling | Cat#:14C10; RRID:AB_561053 |
| Rabbit monoclonal anti-CAND1 | Cell Signaling | Cat#:8759S (D1F2); RRID:AB_11178669 |
| Mouse monoclonal anti-KLHL12 | Cell Signaling | Cat#:9406S (2G2); RRID:AB_2797699 |
| Rabbit monoclonal anti-ubiquityl-PCNA(Lys164) | Cell Signaling | Cat#:13439 RRID:AB_2798219 |
| Mouse monoclonal anti-Ubiquitin (P4D1) | Cell Signaling | Cat#:3936; AB_331292 |
| Rabbit polyclonal anti-KBTBD7 | Abcam | Cat#:ab230126 |
| Mouse monoclonal anti-KLHL4 generated with GenScript | This study | N/A |
| Rabbit monoclonal anti-PAX6 | Cell Signaling | Cat#:60433; RRID:AB_2797599 |
| Rabbit polyclonal anti-FOXG1 | abcam | Cat#:ab18259; RRID:AB_732415 |
| Mouse monoclonal anti-TFAP2A | Novus | Cat#:NB100-74359; RRID:AB_1048155 |
| Rabbit polyclonal anti-TFAP2B | Cell Signaling | Cat#:2509; RRID:AB_2058198 |
| Rabbit monoclonal anti-SOX10 | Cell Signaling | Cat#: 89356; RRID:AB_2792980 |

| | | |
|---|---|---|
| Rabbit monoclonal anti-SLUG/SNAIL2 | Cell Signaling | Cat#:9585; RRID:AB_2239535 |
| Rabbit monoclonal anti-NANOG | Cell Signaling | Cat#:4903; RRID:AB_10559205 |
| Goat polyclonal anti-OCT4 | Santa Cruz Biotechnology, Inc. | Cat#:sc-8628; RRID:AB_653551 |
| Rabbit polyclonal anti-GIT1 | Cell Signaling | Cat#:2919; RRID:AB_2109982 |
| Rabbit polyclonal anti-GIT2 | Cell Signaling | Cat#:6953; RRID:AB_10828712 |
| Rabbit monoclonal Cool2/alpha-PIX | Cell Signaling | Cat#:4573; RRID:AB_2060193 |
| Rabbit polyclonal Cool1/beta-PIX | Cell Signaling | Cat#:4515; RRID:AB_2274365 |
| Mouse monoclonal anti-PAK1,2,3 | Santa Cruz Biotechnology, Inc. | Cat#:sc-166174; RRID:AB_2160693 |
| Rabbit monoclonal anti-CDC42 | Cell Signaling | Cat#:2466; RRID:AB_2078082 |
| Rabbit monoclonal anti-RAC1/2/3 | Cell Signaling | Cat#:2465; RRID:AB_2176152 |
| Rabbit monoclonal anti-RhoA | Cell Signaling | Cat#:2117; RRID:AB_10693922 |
| Mouse monoclonal anti-PAX7 | DSHB | Cat# pax7 RRID:AB_528428 |
| m-IgGk BP-HRP conjugate | Santa Cruz Biotechnology, Inc. | Cat#:sc-516102 |
| HRP Donkey-anti-Mouse IgG | Jackson ImmunoResearch Laboratories | Cat#:715-035-150; RRID:AB_2340770 |
| HRP Donkey-anti-Rabbit IgG | Jackson ImmunoResearch Laboratories | Cat#:711-035-152; RRID:AB_10015282 |

| | | |
|---|---|---|
| HRP Goat-anti-Mouse IgG | Jackson ImmunoResearch Laboratories | Cat#:705-035-147; RRID:AB_2313587 |
| Donkey-anti-Mouse IgG Alexa Fluor 488 | Jackson ImmunoResearch Laboratories | Cat#:715-545-150; RRID:AB_2340846 |
| Donkey-anti-Rabbit IgG Rhodamine Red | Jackson ImmunoResearch Laboratories | Cat#:711-295-152; RRID:AB_2340613 |
| Donkey-anti-Mouse IgG Alexa Fluor 647 | Jackson ImmunoResearch Laboratories | Cat#:715-605-150; RRID:AB_2340862 |
| Donkey-anti-Goat IgG Alexa Fluor 488 | Jackson ImmunoResearch Laboratories | Cat#:705-545-147; RRID:AB_2336933 |
| Normal mouse IgG | Santa Cruz Biotechnology, Inc. | Cat#:Sc-2025; RRID:AB_737182 |
| Bacterial and virus strains | | |
| E.coli: One Shot Stbl3 Chemically competent cells | ThermoFisher Scientific | Cat#:C7373-03 |
| *E. coli*: DH5alpha | ThermoFisher Scientific | Cat#:18265017 |
| *E. coli*: Rosetta2(DE3) | MilliporeSigma | Cat#:71400 |
| Chemicals, peptides, and recombinant proteins | | |
| Polyethylenimine (PEI), Linear, MW 25000, Transfection Grade | Polysciences | Cat#:23966-1 |
| TCEP (Tris(2-carboxyethyl)phosphine hydrochloride)) | Sigma-Aldrich | Cat#:C4706 |
| EGTA | Sigma | Cat#:E3889-25G |
| Dithiothreitol | Invitrogen | Cat#:15508-013 |
| 2-Mercaptoethanol | VWR Life Sciences | Cat#:60-24-2 |
| Urea | FisherChemical | Cat#:U16-3 |
| 16% Formaldehyde solution | Thermo Scientific | Cat#:28908 |
| Doxycycline hyclate | Sigma-Aldrich | Cat#:D9891 |
| Puromycin dihydrochloride | Sigma-Aldrich | Cat#:P8833 |
| G418 disulfate salt solution | Sigma-Aldrich | Cat#:G8168 |
| ML141 (CDC42 inhibitor) | Sigma-Aldrich | Cat#:SML0407 |
| Y-27632 Dihydrochloride (ROCK inhibitor) | Stem Cell Technologies | Cat#:72307 |

| | | |
|---|---|---|
| *S*-Methyl thiomethanesulfonate, MMTS | MilliporeSigma | Cat#:208795 |
| Formic Acid (Optima LC/MS) | Fisher Chemical | Cat#:64-18-6 |
| Trichloroacetic acid | Fisher Chemical | Cat#:A322-100 |
| 3xFLAG peptide | Sigma-Aldrich | Cat#:F4799 |
| $^{6xHIS\text{-}MBP}$GIT1$^{3KD\text{-}R}$ (GIT1-R564D, RK566,567DD) | This study | N/A |
| $^{6xHIS\text{-}MBP}$GIT1 | This study | N/A |
| KLHL4$^{FLAG}$ | This study | N/A |
| **Critical commercial assays** | | |
| Chemiluminescent HRP substrate | Millipore | Cat#:WBKLS0500 |
| Pierce 660nm Protein Assay Reagent | ThermoFisher Scientific | Cat#:22660 |
| **Experimental models: Cell lines** | | |
| Human: iPSC line dCas9-KRAB WTC | Coriell | AICS-0090-391 |
| Human: WA01 hESC H1 line (NIHhESC-10-0043) | Wicell | WAe001-A |
| Human: HEK 293T cells | ATCC | CRL-3216 |
| Human: hTERT RPE-1 cells | ATCC | CRL-4000 |
| **Experimental models: Organisms/strains** | | |
| Fertilized hen eggs from the University of Connecticut | | N/A |
| **Oligonucleotides** | | |
| Silencer® select siRNA control #1 | ThermoFisher Scientific | Cat#:4404021 |
| Silencer® select siRNA control #2 | ThermoFisher Scientific | Cat#:4390846 |
| Silencer® select siRNA KLHL4 #1 | ThermoFisher Scientific | Cat#:4392420 s31872 |
| Silencer® select siRNA KLHL4 #2 | ThermoFisher Scientific | Cat#:4392420 s31873 |
| Silencer® select siRNA GIT1 | ThermoFisher Scientific | Cat#:4390824 s26308 |
| Silencer® select siRNA GIT2 | ThermoFisher Scientific | Cat#:4392420 s18975 |
| Silencer® select siRNA alpha-PIX | ThermoFisher Scientific | Cat#:4392420 s18123 |
| Silencer® select siRNA beta-PIX | ThermoFisher Scientific | Cat#:4392420 s16949 |
| Silencer® select siRNA KLHL5 | ThermoFisher Scientific | Cat#:4392420 s27396 |
| Silencer® select siRNA PAK1 | ThermoFisher Scientific | Cat#:4390824 s10021 |

| | | |
|---|---|---|
| Silencer® select siRNA PAK2 | ThermoFisher Scientific | Cat#:4390824 s10021 |
| Silencer® select siRNA PAK3 | ThermoFisher Scientific | Cat#:4390824 s534740 |
| control guide: GGACTAAGCGCAAGCACCTA | This study | N/A |
| KLHL4 guide: TATGAGCATAGAGGGACCC | This study | N/A |
| HCR probe: Myc-N (gallus gallus) | (Lignell et al., 2017) | N/A |
| HCR probe: Msx1 (gallus gallus), Split initiator | Molecular Instruments | N/A |
| HCR probe: Dlx5 (gallus gallus), Split initiator | Molecular Instruments | N/A |
| qPCR primers | Table S5 | N/A |
| **Recombinant DNA** | | |
| Plasmid: B9-sgKLHL4 | This study | N/A |
| Plasmid: B9-sgControl | This study | N/A |
| Plasmid: Mission® pLKO1-puro-shGIT1 | Sigma | TRCN0000436470 |
| Plasmid: Mission® pLKO1-puro-shGIT2 | Sigma | TRCN0000364545 |
| Plasmid: pInducer20-FLAG-CUL3 | This study | N/A |
| Plasmid: pInducer20-FLAG-CUL3-S53N | This study | N/A |
| Plasmid: pInducer20-FLAG-CUL3-Y62F | This study | N/A |
| Plasmid: pInducer20-FLAG-CUL3-I86V | This study | N/A |
| Plasmid: pInducer20-FLAG-CUL3-I276V | This study | N/A |
| Plasmid: pInducer20-FLAG-CUL3-V285A | This study | N/A |
| Plasmid: pInducer20-FLAG-CUL3-E752Q | This study | N/A |
| Plasmid: pInducer20-KLHL4-WT-CRiQ5-3xFLAG | This study | N/A |
| Plasmid: pInducer20-KLHL4-I287V-CRiQ5-3xFLAG | This study | N/A |
| Plasmid: pInducer20-KLHL4-AP1-CRiQ5-3xFLAG | This study | N/A |
| Plasmid: pInducer20-HA-PAK1 | This study | N/A |
| Plasmid: pInducer20-HA-PAK1-Ub | This study | N/A |
| Plasmid: pInducer20-HA-PAK1-UbI44A | This study | N/A |
| Plasmid: pCDNA5 KLHL4-3xFLAG | This study | N/A |
| Plasmid: pCDNA5 KLHL4-K330E-3xFLAG | This study | N/A |
| Plasmid: pCDNA5 KLHL4-I287V-3xFLAG | This study | N/A |
| Plasmid: pCDNA5 KLHL4-AP1-(DDED475,544,546,591RRRR)-3xFLAG | This study | N/A |
| Plasmid: pCDNA5 KLHL4-AP2-(DD644,648RR)-3xFLAG | This study | N/A |
| Plasmid: pCDNA5 KLHL4-AP3-(EE699,700RR)-3xFLAG | This study | N/A |
| Plasmid: pCDNA5 KLHL4ΔCUL3(Y206A)-3xFLAG | This study | N/A |
| Plasmid: pCDNA5-KLHL4-82-Cterm-3xFLAG | This study | N/A |
| Plasmid: pCDNA5-KLHL4-128-Cterm-3xFLAG | This study | N/A |
| Plasmid: pCDNA5 KLHL5-3xFLAG | This study | N/A |
| Plasmid: pCDNA5 KLHL5-I377T-3xFLAG | This study | N/A |
| Plasmid: pCS2-KLHL5-3xHA | This study | N/A |
| plasmid: pCS2-KLHL17-3xFLAG | This study | N/A |
| plasmid: pCS2-KLHL17-L239P-3xFLAG | This study | N/A |
| plasmid: pCS2-KLHL17-P234L-3xFLAG | This study | N/A |
| plasmid: pCS2-KLHL17-3xHA | This study | N/A |

| | | |
|---|---|---|
| plasmid: pCS2-KLHL11-3xFLAG | This study | N/A |
| plasmid: pCS2-KLHL11-T243S-3xFLAG | This study | N/A |
| plasmid: pCS2-KLHL11-3xHA | This study | N/A |
| plasmid: pCDNA5-KBTBD8-3xFLAG | [35] | N/A |
| plasmid: pCDNA5-KBTBD8 W220C-3xFLAG | This study | N/A |
| plasmid: pCDNA5-KBTBD8-3xHA | [35] | N/A |
| plasmid: pCDNA5 KLHL12-3xFLAG | [36] | N/A |
| plasmid: pCS2-KLHL12-HA | [36] | N/A |
| plasmid: pCDNA5-KLHL12 L227P-3xFLAG | This study | N/A |
| plasmid: pCS2-KLHL18-3xFLAG | This study | N/A |
| plasmid: pCS2-KLHL18-3xHA | This study | N/A |
| plasmid: pCS2-KLHL18-D244G-3xFLAG | This study | N/A |
| plasmid: pCS2-KLHL41-3xFLAG | This study | N/A |
| plasmid: pCS2-KLHL41-3xHA | This study | N/A |
| plasmid: pCS2-KLHL41D240G-3xFLAG | This study | N/A |
| plasmid: pCS2-KLHL20-3xFLAG | This study | N/A |
| plasmid: pCS2-KLHL20-3xHA | This study | N/A |
| plasmid: pCS2-KLHL20-R266C-3xFLAG | This study | N/A |
| plasmid: pCS2-KBTBD4-3xFLAG | This study | N/A |
| plasmid: pCS2-KBTBD4-3xHA | This study | N/A |
| plasmid: pCS2-KBTBD4-R220G-3xFLAG | This study | N/A |
| plasmid: pCS2-KLHL8-3xFLAG | This study | N/A |
| plasmid: pCS2-KLHL8-3xHA | This study | N/A |
| plasmid: pCS2-KLHL8-C87F-3xFLAG | This study | N/A |
| plasmid: pCDNA5-KLHL1-3xFLAG | This study | N/A |
| plasmid: pCS2-KLHL11-3xHA | This study | N/A |
| plasmid: pCDNA5 KLHL1-L305P-3xFLAG | This study | N/A |
| plasmid: pHAGE-puro-KLHL36-FLAG-HA | This study | N/A |
| plasmid: pHAGE-puro-KLHL36-T288M-FLAG-HA | This study | N/A |
| plasmid: pCS2-3xHA-GIT1 | This study | N/A |
| plasmid: pCS2-3xHA-GIT1-1-620 | This study | N/A |
| plasmid: pCS2-3xHA-GIT1-1-420 | This study | N/A |
| plasmid: pCS2-3xHA-GIT1-1-396 | This study | N/A |
| plasmid: pCS2-3xHA-GIT1-373-629 | This study | N/A |
| plasmid: pCS2-3xHA-GIT1-429-629 | This study | N/A |
| plasmid: pCS2-3xHA-GIT1-429-541 | This study | N/A |
| plasmid: pCS2-3xHA-GIT1-429-580 | This study | N/A |
| plasmid: pCS2-3xHA-GIT1-429-518 | This study | N/A |
| plasmid: pCS2-3xHA-GIT1-429-490 | This study | N/A |
| plasmid: pCS2-3xHA-GIT1-429-567 | This study | N/A |
| plasmid: pCS2-3xHA-GIT1-429-560 | This study | N/A |
| plasmid: pCS2-HA3x-GIT1-429-551 | This study | N/A |
| plasmid: pCS2-HA3x-GIT1-Δ560-570 | This study | N/A |
| plasmid: pCS2-3xHA-GIT1-Y563F | This study | N/A |
| plasmid: pCS2-3xHA-GIT1-Y563D | This study | N/A |
| plasmid: pCS2-3xHA-GIT1-RK3A (R564A,RK566AA) | This study | N/A |
| plasmid: pCS2-3xHA-GIT1-RK3D (R564D,RK566DD) | This study | N/A |
| plasmid: pCS2-3xHA-PAK1 | This study | N/A |
| plasmid: pCS2-3xHA-PAK1-ubiquitin | This study | N/A |

| | | |
|---|---|---|
| plasmid: pCS2-3xHA-PAK1-ubiquitin-I44A | This study | N/A |
| plasmid: pCS2-3xHA-PAK1ΔPIX(PRP192-194AAA) | This study | N/A |
| plasmid: pCS2-3xHA-PAK2 | This study | N/A |
| plasmid: pCS2-3xHA-PAK3 | This study | N/A |
| plasmid: pCMV-FLAG3x-PAK1 | This study | N/A |
| plasmid: CMV-FLAG3x-PAK2 | This study | N/A |
| plasmid: pCMV-FLAG3x-PAK3 | This study | N/A |
| plasmid: pCS2-MYC-PAK1 | This study | N/A |
| plasmid: pCS2-MYC-PAK2 | This study | N/A |
| plasmid: pCS2-MYC-PAK3 | This study | N/A |
| plasmid: pCS2-His-Ubiquitin | [36] | N/A |
| plasmid: pCS2-His-Ubiquitin-K0 (all K residues mutated to R) | [36] | N/A |
| plasmid: pCMV-3xFLAG-GFP-CDC42-T17N | This study | N/A |
| plasmid: pET28a-His-MBP-GIT1 | This study | N/A |
| plasmid: pET28a-His-MBP-GIT1-RK3D (R564D,RK566DD) | This study | N/A |
| plasmid: pCS2-HA3x-α-PIX-SH3*(WW171,172PG) | This study | N/A |
| plasmid: pCS2-HA3x-β-PIX-SH3*(WW197,198PG) | This study | N/A |
| plasmid: pCS2-MYC-β-PIX | This study | N/A |
| Software and algorithms | | |
| Fiji | (Schindelin et al. 2012) | RRID:SCR_002285 |
| CompPASS | (Sowa et al., 2009) | N/A |
| ImageJ | N/A | PRID:SCR_003070 |
| Prism | GraphPad | http://www.graphpad.com/ |
| Other | | |
| Bovine Serum Albumin | Sigma | Cat#:A8806-5G |
| Lipofectamine RNAiMAX | ThermoFisher Scientific | Cat#:13778030 |
| Complete, EDTA-free protease inhibitor cocktail tablets from Roche | Sigma-Aldrich | Cat#:11873580001 |
| Hoechst 33342 | ThermoFisher Scientific | Cat#:H3570 |
| Opti-MEM (1X) Reduced Serum Medium | GIBCO | Cat#:31985-070 |
| DMEM (1X) +GlutaMAX™-I | GIBCO | Cat#:51985091 |
| Trypsin-EDTA (0.25%) | GIBCO | Cat#:25200056 |
| Fetal Bovine Serum | VWR LifeScience | Cat#:97068-107 |
| Ni-NTA | QIAGEN | Cat#:30210 |
| Lipofectamine RNAiMAX | ThermoFisher | Cat#:13778150 |
| ANTI-FLAG® M2 Affinity Agarose Gel slurry | Sigma-Aldrich | Cat#:CA2220 |

| | | |
|---|---|---|
| Protein G-Agarose | Sigma-Aldrich | Cat#:11243233001 |
| Matrigel hESC-Qualified Matrix | Corning | Cat#:354277 |
| mTeSR$^{TM}$1 | StemCell Technologies Inc. | Cat#:05871/05852 |
| STEMdiff$^{TM}$ Neural Induction Medium | StemCell Technologies Inc. | Cat#:05831 |
| APEL2 medium | StemCell Technologies Inc. | Cat#:05270 |
| Collagenase | StemCell Technologies Inc. | Cat#:07909 |
| Acctuase | StemCell Technologies Inc. | Cat#:07920 |
| Anti Dig-AP | Roche Diagnostics GmbH | Cat#:11093274910 |
| NBT (4-Nitro blue tetrazolium chloride, solution) | Roche Diagnostics GmbH | Cat#:11383213001 |
| BCIP (4-toluidine salt, solution) | Roche Diagnostics GmbH | Cat#:11383221001 |
| Invitrogen ProLong$^{TM}$ Gold Antifade Mountant | Fisher Scientific | Cat#: P36934 |
